# Reinforced identity confers cells with cancer hallmarks

**DOI:** 10.1101/2024.11.01.621626

**Authors:** Haixia Yang, Pengqi Wang, Pengcheng Tan, Qiaomei Xue, Hua Wu, Kezhang He, Bowen Wang, Dan Wang, Tianhua Ma, Sheng Ding

## Abstract

Genetic alterations trigger tumorigenesis in a cell type-specific manner, but the biological basis for this, remains unclear. Here we define reinforced original cell identity established by interaction between genetic alterations and original cell type is a pivotal determinant of tumor development. Using AML-M5^MLL-AF9^ as a model system, we demonstrated that MLL-AF9 enhanced key transcriptional factors (TFs) expression of promonocyte and cooperated with them to amplify their downstream programs for AML hallmarks. Interactions between MLL-AF9 and promonocyte endowed cancer cells with a reinforced promonocyte identity. Reinforced intrinsic cell identity also sustained cancer hallmarks across cancer types. Furthermore, we identified narciclasine, as an effective disruptor of reinforced promonocyte identity, effectively treating AML-M5^MLL-AF9^. This study not only identified reinforced cell identity as a fundamental tumor driver beyond genetic mutations but also offers a proof-of-concept for reinforced identity- targeted cancer therapy.

## Introduction

High-throughput sequencing studies have shown pronounced tissue specificity in cancer driver mutations (*1, 2*). Subsequent studies have underscored that several oncogenic mutations possess a cell type-dependent transformative potential. (*3–6*) (*3, 7, 8*). However, the biological basis for this, remains unclear.

The oncogenic MLL-AF9 fusion, one of the most frequent MLL/HRX/ALL-1 rearrangements resulting from t(9;11) (p22;q23) translocation, is exclusively found in monocytic acute myeloid leukemia (AML-M5) and infantile acute lymphocytic leukemia (ALL) patients (*9, 10*). The expression of MLL-AF9 fusion leads to the development of acute leukemia, specifically AML and ALL, but not other types of cancers (*11, 12*). Patient-derived AML-M5^MLL-AF9^ cells distinctly exhibited promonocyte identity (*13*). Hence, we set out to use AML-M5^MLL-AF9^ as a model system to investigate how interaction between genetic alterations and specific cell type influence tumor development.

## Results

### AML-M5^MLL-AF9^ cells depend on reinforced promonocyte identity

We analyzed two AML-M5^MLL-AF9^ cell lines (THP1 and MOLM-13 cells), which exhibited the typical morphology of promonocytes, observed in other AML-M5^MLL-AF9^ cells [11-12]. Transcriptomic comparison revealed both THP1 and MOLM-13 cells share similar gene expression profiles with promonocytes (fig. S1A), aligning with previous single-cell RNA sequencing (scRNA-seq) analysis of cancer cells from AML-M5 patients (*13*). A set of TFs essential for promonocyte identity, including IRF8, SPI1, GFI1 and CEBPA, were indeed highly expressed in both THP1 and MOLM-13 cells (Fig.1A). The promoters of these TFs exhibited transcriptionally active marks, the enrichment of both H3K27ac and H3K4me3 (fig. S1B). In addition, within the open chromatin regions identified using ATAC-seq data from AML-M5^MLL- AF9^ patients, DNA binding motif analysis revealed a significant enrichment of specific binding sites for IRF8, SPI1 and GFI1 (Fig. 1B). These observations collectively confirmed the promonocyte identity of AML-M5^MLL-AF9^ cells. Furthermore, we compared the expression of key promonocyte TFs, such as IRF8, SPI1, GFI1, CEBPA and MYB, between AML-M5^MLL-AF9^ cells and normal promonocytes. Intriguingly, all these TFs exhibited significantly higher expression in AML-M5^MLL-AF9^ than in normal promonocyte (Fig. 1C), indicating a reinforced promonocyte identity in AML-M5^MLL-AF9^ cells.

**Fig. 1.**
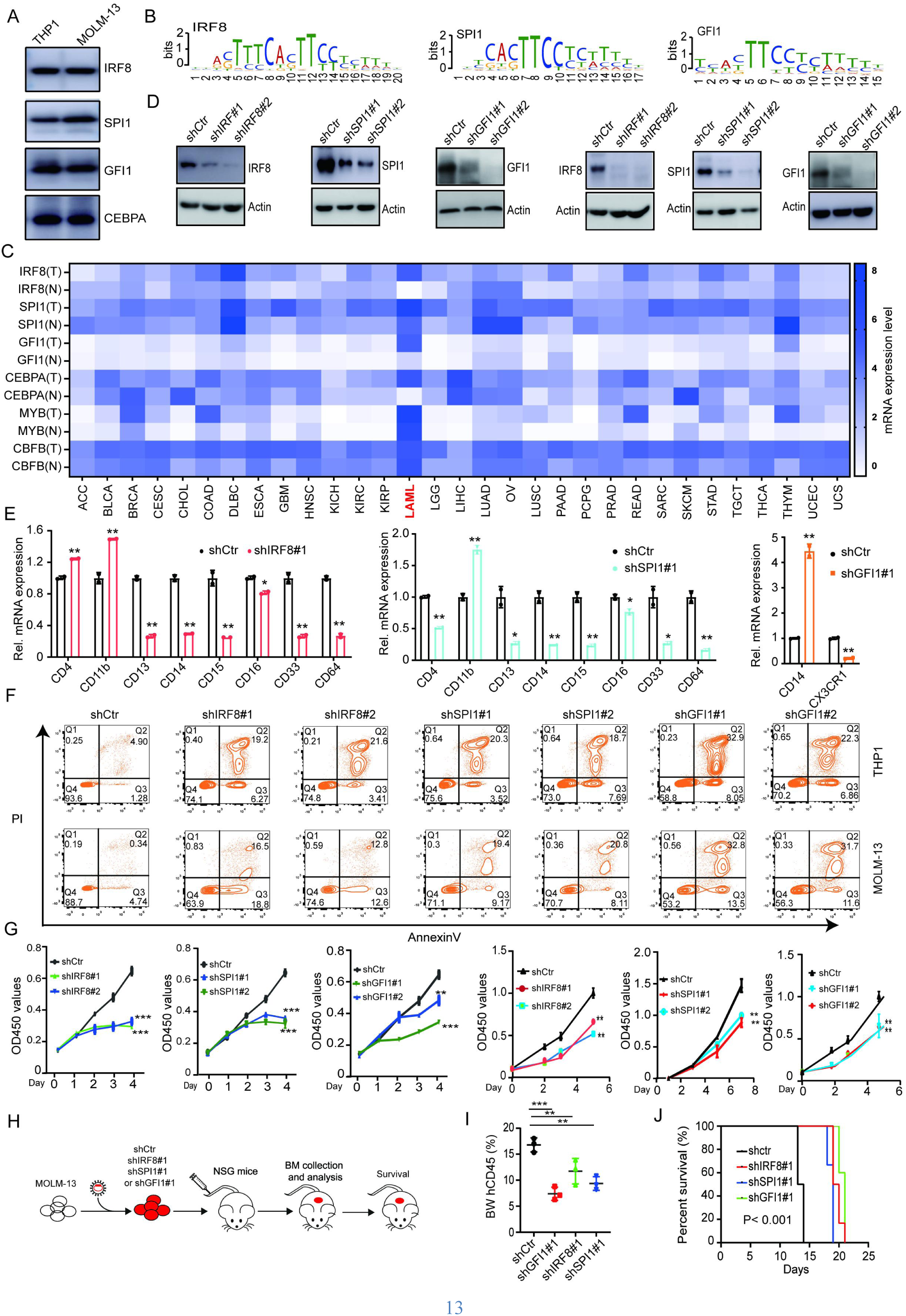
AML-M5MLL-AF9 cells depend on reinforced promonocyte identity (**A**) Western blot analysis showing the expression levels of key promonocyte TFs in THP1 and MOLM-13 cells. (**B**) De novo motif analysis conducted with MEME on sequences from open chromatin regions identified by ATAC-seq in THP1 and MOLM-13 cells. (**C**) Heatmap comparison of relative mRNA expression levels of key promonocyte TFs between various indicated cancer types and their normal tissue counterparts. ACC, Adrenocortical carcinoma; BLCA, Bladder Urothelial Carcinoma; BRCA, Breast invasive carcinoma; CESC, Cervical squamous cell carcinoma and endocervical adenocarcinoma; CHOL, Cholangiocarcinoma; COAD, Colon adenocarcinoma; DLBC, Lymphoid Neoplasm Diffuse Large B-cell Lymphoma; ESCA, Esophageal carcinoma; GBM, Glioblastoma multiforme; HNSC, Head and Neck squamous cell carcinoma; KICH, Kidney Chromophobe; KIRC, Kidney renal clear cell carcinoma; KIRP, Kidney renal papillary cell carcinoma; LAML, Acute Myeloid Leukemia; LGG, Brain Lower Grade Glioma; LIHC, Liver hepatocellular carcinoma; LUAD, Lung adenocarcinoma; OV, Ovarian serous cystadenocarcinoma; LUSC, Lung squamous cell carcinoma; PAAD, Pancreatic adenocarcinoma; PCPG, Pheochromocytoma and Paraganglioma; PRAD,Prostate adenocarcinoma; READ, Rectum adenocarcinoma; SARC, Sarcoma; SKCM, Skin Cutaneous Melanoma; STAD, Stomach adenocarcinoma; TGCT, Testicular Germ Cell Tumors; THCA, Thyroid carcinoma; THYM, Thymoma; UCEC, Uterine Corpus Endometrial Carcinoma; UCS, Uterine Carcinosarcoma. (**D**) Western blot (bottom) results validating specified shRNA-mediated knockdown of IRF8, SPI1, and GFI1 in THP1 (a) and MOLM-13 (b) cells, with scramble shRNA (shCtr) serving as a control. (**E**) RT-qPCR analysis indicating changes in the expression of indicated promonocyte surface markers in THP1 cells upon knockdown of IRF8, SPI1, or GFI1 using specified shRNA. (**F**) Apoptosis in THP1 and MOLM- 13 cells transduced with indicated shRNAs for 72h. (**G**) Cell viability of THP1 and MOLM-13 cells transduced with indicated shRNAs for 72h. (**H**) Schematic workflow of *in-vivo* transplantation of MOLM-13 cells infected with shCtr, shIRF8#1, shSPI1#1 or shGFI1#1. (**I**) Flow cytometry analysis showing the percentage of human CD45^+^ leukemia cells in the bone marrow of NSG recipient mice 7 days after transplantation with MOLM-13 cells transduced with the specified shRNAs. (**J**) Survival curve of NSG mice following transplantation as described in (H).

To access the importance of reinforced promonocyte identity in AML-M5^MLL-AF9^ cells, we individually knocked down three key promonocyte TFs (IRF8, SPI1 and GFI1) in THP1 and MOLM-13 cells (Fig.1D and fig. S1C). As expected, the knockdown of IRF8, SPI1 or GIF1 significantly downregulated most promonocyte surface markers in both cell lines (Fig. 1E and fig. S1D), disrupting the reinforced promonocyte identity. Moreover, this knockdown led to pronounced apoptosis and cell-cycle arrest at the S-phase in both THP1 and MOLM-13 cells (Fig. 1F and fig. S1E), resulting in a significant reduction in their proliferation and clonogenic activity (Fig. 1G and fig. S1F). When intravenously injecting MOLM-13 cells and their counterparts with IRF8, SPI1 or GFI1 knockdown, into NOD scid gamma (NSG) immunodeficient mice (Fig. 1H), we found these knockdown significantly decreased the presence of AML-M5^MLL-AF9^ cells in bone marrow (Fig. 1I), and markedly prolonged the survival of NSG mice injected with AML-M5^MLL- AF9^ cells (Fig. 1J). These results collectively indicated that AML-M5^MLL-AF9^ cells depend on a reinforced promonocyte identity to maintain their cancer hallmarks.

### Reinforced promonocyte identity endows AML-M5^MLL-AF9^ cells with cancer hallmarks

*MLL-AF9* fusion is a driver for AML-M5^MLL-AF9^ (*14–20*). Knockdown of MLL-AF9 led to pronounced apoptosis and cell-cycle arrest at the S phase in both THP1 and MOLM-13 cells, suppressing their proliferation and clonogenic activity (fig. S2, A to D). These effects paralleled those observed when disrupting their reinforced promonocyte identity, indicating a possible relationship between MLL-AF9 and reinforced promonocyte identity. Intriguingly, MLL-AF9 knockdown markedly reduced the expression of key promonocyte TFs (IRF8, SPI1, CEBPA, GFI1, RUNX1, RUNX2, MYB, and LMO2) and most promonocyte surface markers in both cell lines (Fig. 2A and fig. S2E). These findings indicated that MLL-AF9 is essential for the reinforced promonocyte identity in AML-M5^MLL-AF9^ cells.

**Fig. 2.**
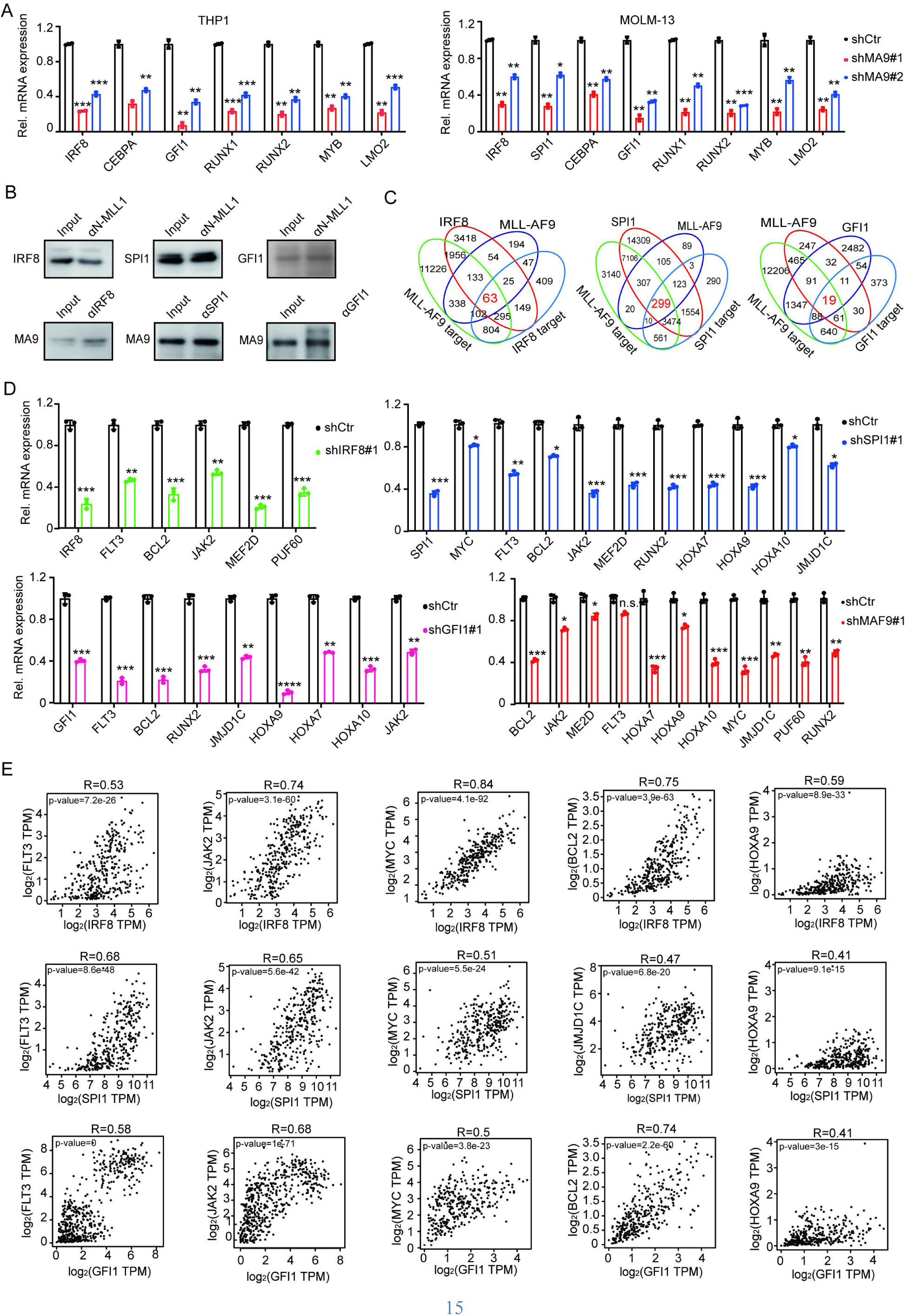
Reinforced promonocyte identity endows AML-M5^MLL-AF9^ cells with cancer hallmarks (**A**) RT-qPCR analysis of the expression levels of indicated promonocyte key TFs in THP1 (left) and MOLM-13 (right) cells following transduction with scramble (shCtr) and MLL-AF9 (shMA9#1 and shMA9#2) shRNAs for 72h. (**B**) Endogenous co-immunoprecipitation analysis illustrating MLL-AF9’s interaction with IRF8, SPI1, and GFI1. (**C**) Venn diagrams indicating the overlap between genes bound by the indicated proteins and that downregulated following specified gene knockdown. The number of common downregulated and co-bound genes between MLL-AF9 and IRF8, SPI1 or GFI1 was highlighted in red. (**D**) RT-qPCR analysis revealing genes commonly downregulated in THP1 cells by knockdown of MLL-AF9 and TFs IRF8, SPI1 or GFI1. (**E**) Dot plots illustrating the positive expression correlation between IRF8, SPI1, or GFI1 and specified genes in AML patients’ samples, as analyzed using GEPIA 2.0.

To investigate the underling mechanism, we mapped genomic binding of MLL-AF9 in THP1 and MOLM-13 cells using Chromatin Immunoprecipitation Sequencing (ChIP-seq), considering that it can bind to genome (*21, 22*). ChIP-seq analysis identified 4631 and 1362 MLL-AF9 binding peaks in THP1 and MOLM-13 cells, respectively, with 1027 peaks common to both (fig. S2, F and G), particularly including active regulatory regions of key promonocyte TFs like IRF8, SPI1, GFI1, CEBPA, MYB, RUNX1, RUNX2 and BMI1 (fig. S3). This led to the hypothesis that MLL-AF9 could promote the expression of key promonocyte TFs through directly binding to their promoters/enhancers. To test this hypothesis, we inserted various 1.5-2kb DNA fragments which span 4.5∼6 kb promoters/enhancers of IRF8, SPI1, GFI1 and CEBPA into a luciferase-based reporter system, and measured reporter expression in 293T cells ectopically expressing MLL-AF9. We found that MLL-AF9 can significantly increase the reporter gene expression via at least one regulatory sequence from the promoter/enahcner of each key promonocyte TF (fig. S2H), confirming these regions can be bound by MLL-AF9 for activation of direct downstream genes. The protein expression of IRF8, SPI1, GFI1, and CEBPA in THP1 and MOLM-13 cells were consistently downregulated following MLL-AF9 knockdown (fig. S2I). Thus, our results collectively demonstrated that MLL-AF9 promoted the expression of key promonocyte TFs through directly binding to their promoters/enhancers in AML-M5^MLL-AF9^ cells.

Recognizing a significant portion of MLL-AF9 binding peaks are located within other regions besides key promonocyte TFs, we performed DNA binding motif analysis for these regions. This revealed these motifs that can be bound by key promonocyte TFs, such as IRF8, SPI1, GFI1, CEBPA, MYB, RUNX1 and FLI1 (fig. S4A). Subsequently, analysis of chromatin binding of IFR8, SPI1 and GFI1 by ChIP-seq, showing that 82.74% of MLL-AF9 binding peaks in THP1 cells overlapped with binding peaks of these TFs, while in MOLM-13 cells, 76.5% of MLL-AF9 binding peaks coincided with these TFs’ binding peaks (fig. S4, B and F). Consistently, MLL- AF9 interacted with IRF8, SPI1 or GFI1 (Fig. 2B). The domains of MLL-AF9 responsible for these interactions were SNL and CxxC domains for IRF8, CxxC for SPI1, and CxxC for GFI1 (fig. S4, G to I).

To determine whether MLL-AF9 and key promonocyte TFs could cooperatively regulate the transcription in their co-bound regions, we performed comparative RNA-seq analysis in THP1 and MOLM-13 cells following knockdown of MLL-AF9, IRF8, SPI1, or GFI1. Among genes co-bound by MLL-AF9 and IRF8, 13.2% (63 genes) in THP1 cells and 6.9% (169 genes) in MOLM13 cells were commonly downregulated after knocking down either MLL-AF9 or IRF8. For genes co-bound by MLL-AF9 and SPI1, the percentage of commonly downregulated genes were 35.8% (299 genes) in THP1 cells and 5.6% (435 genes) in MOLM13 cells. Similarly, for MLL-AF9 and GFI1, the common downregulation affected 12.4% (19 genes) in THP1 cells (Fig. 2C and fig. S6A). GO analysis of these co-regulated genes highlighted their roles in biological process essential for promonocyte function, including transcriptional regulation, hematopoiesis, differentiation, cell proliferation and anti-apoptosis (fig. S5). Notably, a subset of these co-regulated genes, which are essential for promonocyte traits such as cell growth (*FLT3, JAK2, MEF2D, CDK6* and *MYC*), maintenance of undifferentiated state (*HOXA* family, *JMJD1C*, *MEIS1*, *RUNX1* and *RUNX2*) and anti-apoptosis (*BCL2*) (Fig. 2D and fig. S6B) are key for AML hallmarks [25-30]. Consistently, protein levels of these genes in both THP1 and MOLM-13 cells were decreased following knockdown of IRF8, SPI1, GFI1 or MLL-AF9 (fig. S6, C and D). IRF8, SPI1, and GFI1 each exerted differential regulation of these genes, suggesting their distinct roles in promoting promonocyte identity and AML hallmarks, warranting further investigations. Importantly, there was a positive expression correlation between key promonocyte TFs and their downstream genes key for AML hallmarks in AML-M5^MLL-AF9^ patients’ samples (Fig. 2E and fig. S6E).

In summary, our studies not only showed MLL-AF9 enhances the expression of key promonocyte TFs, but also revealed that MLL-AF9 collaborates with these TFs to amplify the expression of TFs’ downstream genes, particularly those key for AML hallmarks. Consequently, this reinforced promonocyte identity in AML-M5^MLL-AF9^ cells drives manifestation of cancer hallmarks.

### Reinforced promonocyte identity enable genome-wide chromatin binding of MLL-AF9

The widespread genomic co-occupancy of key promonocyte TFs alongside MLL-AF9 suggests that these TFs may facilitate MLL-AF9’s binding to the genome. To examine this hypothesis, we analyzed ATAC-seq data from THP1 and MOLM-13 cells following the knockout of IRF8, as well as the knockdown of SPI1 or GFI1. Remarkably, the knockout of IRF8, along with the knockdown of SPI1 or GFI1, resulted in a significant reduction in genome-wide chromatin accessibility in THP1 cells, affecting 61.2%, 56.1%, and 93.1% of initially accessible loci respectively (Fig. 3, A and B; fig. S7A). Similarly, in MOLM-13 cells, knockdown of SPI1 or GFI1 led to a substantial reduction in chromatin accessibility, impacting 76.8% and 70% of initially accessible loci (Fig. 3, C and D; fig. S7B). Expect for MOLM-13 cells with SPI1 knockdown, total number of chromatin accessible loci in other KO/KD cells were dramatically reduced. These results indicated that KO/KD of key promonocyte TFs resulted in broadly suppressed chromatin accessibility, which may lead to disruption of MLL-AF9’s genome-wide chromatin binding. To confirm this, we performed ChIP-seq in THP1 cells and MOLM-13 cells, observing a substantial decrease in the MLL-AF9 chromatin occupancy following knockout of IRF8, or knockdown of SPI1 or GFI1 (Fig. 3, E and F; fig. S7, C and D). Specifically, in THP1 cells with knockout of IRF8, or knockdown of SPI1 or GFI1, we observed an almost complete loss of both chromatin accessibility and MLL-AF9 occupancy at regions responsible for transcriptional regulation of genes key for AML hallmarks (HOXA family, JMJD1C, LMO2, RUNX1, RUNX2, ZEB2, MESI1, MYC, FLT3, and BCL2) (Fig. 3G and fig. S7E).

**Fig. 3.**
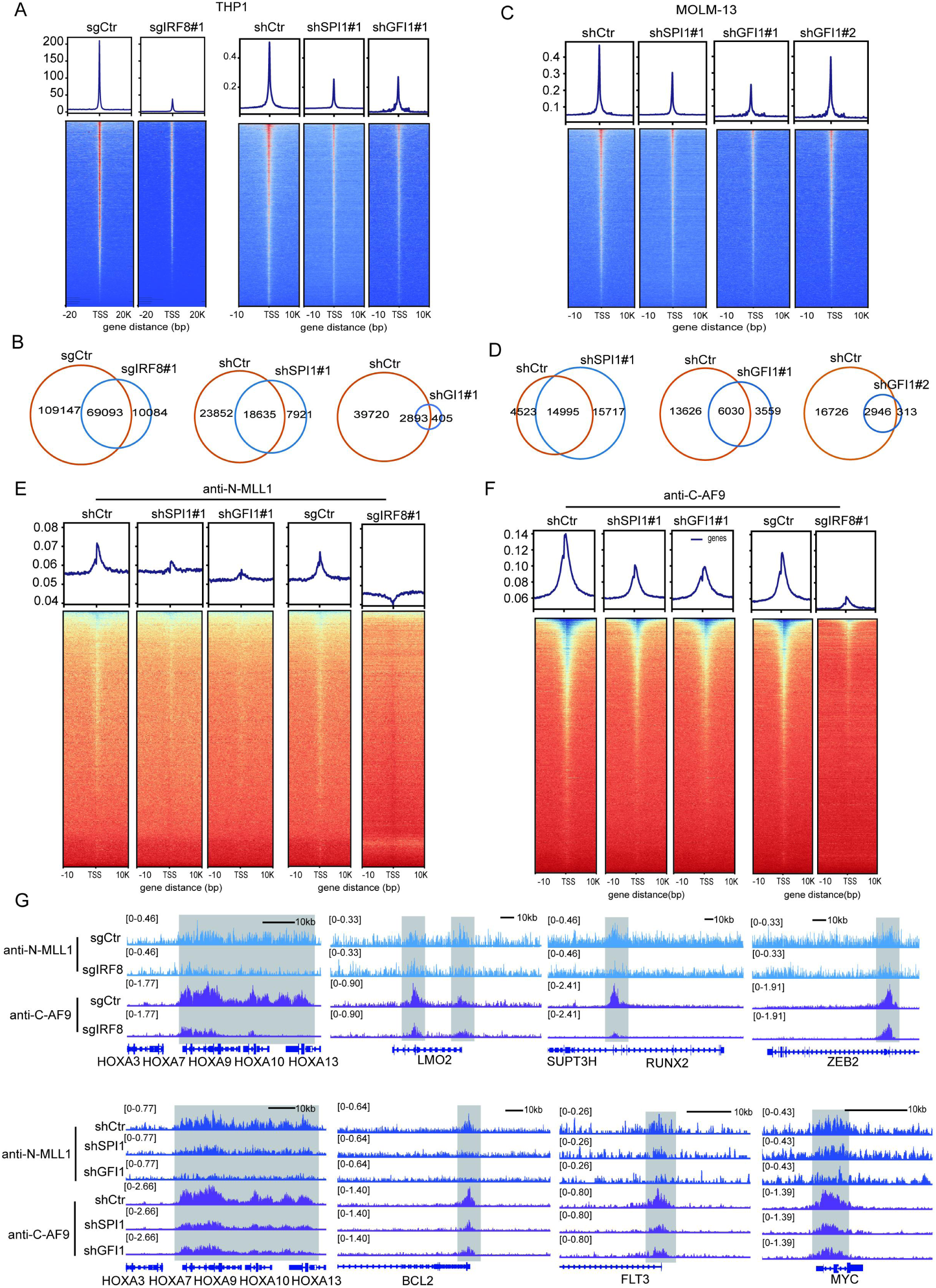
Reinforced promonocyte identity enable genome-wide chromatin binding of MLL-AF9 (**A**) Metaplots (top) and density plots (bottom) illustrating ATAC-seq profiles in THP1 cells post-transduction with control (sgCtr/shCtr) or indicated sgRNAs/shRNAs. (**B**) Venn diagrams illustrating the overlap of ATAC-seq peaks in THP1 cells as described in (A). (**C**) Metaplots (top) and density plots (bottom) showing ATAC-seq profiles in MOLM-13 cells after transduction with control (sgCtr/shCtr) or specified sgRNAs/shRNAs. (**D**) Venn diagrams depicting the overlap of ATAC-seq peaks in MOLM-13 cells as shown in (C). (**E**) Metaplots (top) and density plots (bottom) presenting ChIP-seq profiles for N-MLL1 (N-terminal of MLL1) in THP1 cells following transduction with control (sgCtr/shCtr) or indicated sgRNAs/shRNAs. (**F**), Metaplot (top) and density plot (bottom) displaying ChIP-seq profiles for C-AF9 (C-terminal of AF9) in THP1 cells after transduction with control or specified sgRNAs/shRNAs. (**G**) ChIP-seq gene tracks revealing MLL-AF9 binding peaks at specified gene loci in THP1 cells following transduction with control or indicated sgRNAs/shRNAs.

In summary, highly expressed key promonocyte TFs are instrumental in ensuring chromatin remains accessible within AML-M5^MLL-AF9^ cells. This structural configuration enables the extensive genomic co-occupancy by oncogenic MLL-AF9, which in turn significantly boosts the transcription of adjacent genes, including genes for AML hallmarks. The culmination of these effects is establishment of a reinforced promonocyte identity that defines AML hallmarks in these cells.

### Reinforced cell identity confers cancer hallmarks in various cancer types

To determine whether reinforced cell identity driving cancer features is also applied in various cancer types, we firstly analyzed vulnerabilities across cancer types. Our analysis revealed that in various cancer types, including small cell lung cancer (SCLC), most breast carcinoma subtypes (excluding triple negative breast cancer), melanoma, astrocytoma, glioblastoma, prostate adenocarcinoma, liver cancer, colorectal cancer, esophageal adenocarcinoma, multiple myeloma, acute lymphoblastic leukemia (ALL), and acute myeloid leukemia (AML), cell viability was significantly impaired by the knockout of their corresponding identity-associated transcription factors (TFs) (Fig. 4A and fig. S8A). In the case of AML classified into eight subtypes (M0 to M7) under the FAB classification, we found that all AML cell lines from M2 to M7 subtypes (AML-M0 and M1 were excluded from analysis due to limited data availability) documented in DepMap exhibited high dependency on the expression of their respective identity-associated key TFs (Fig. 4B). These results suggest that various cancer types depends on the original cell identity.

**Fig. 4.**
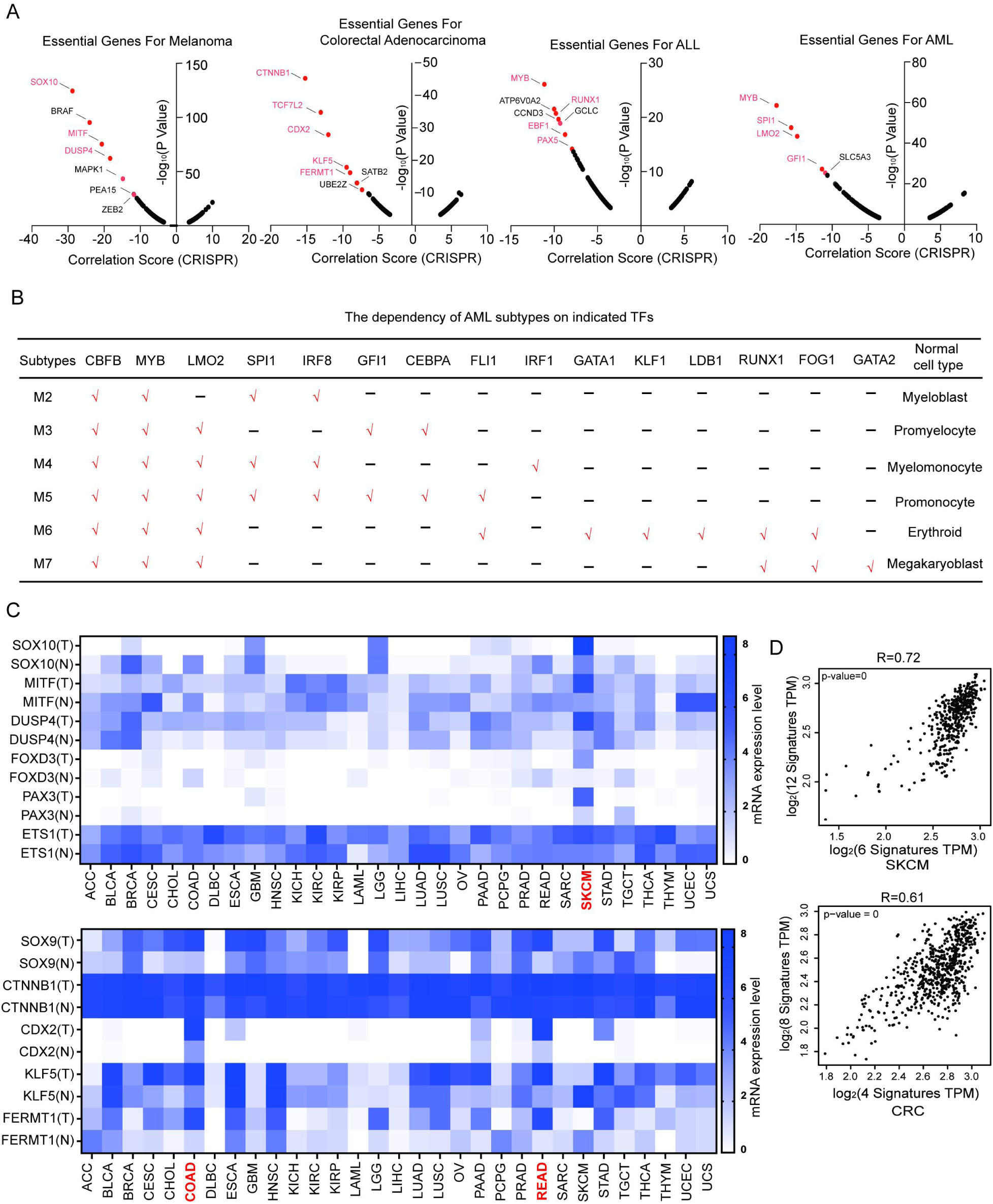
Reinforced cell identity confers cancer hallmarks in various cancer types (**A**) Graphic representations depicting the correlation between the knockout of specific genes and the survival of multiple cancer types. Identity genes were highlighted in red. (**B**) List of key transcription factors essential for the maintenance of AML subtypes M2-M7, and their roles as identity genes in indicated hematological lineages. (**C**) Heatmap showing the relative expression levels of key TFs that define the identity of neural crest cell (top) and intestinal stem cell identity (bottom), comparing various cancer types to their normal tissue counterparts. ACC, Adrenocortical carcinoma; BLCA, Bladder Urothelial Carcinoma; BRCA, Breast invasive carcinoma; CESC, Cervical squamous cell carcinoma and endocervical adenocarcinoma; CHOL, Cholangiocarcinoma; COAD, Colon adenocarcinoma; DLBC, Lymphoid Neoplasm Diffuse Large B-cell Lymphoma; ESCA, Esophageal carcinoma; GBM, Glioblastoma multiforme; HNSC, Head and Neck squamous cell carcinoma; KICH, Kidney Chromophobe; KIRC, Kidney renal clear cell carcinoma; KIRP, Kidney renal papillary cell carcinoma; LAML, Acute Myeloid Leukemia; LGG, Brain Lower Grade Glioma; LIHC, Liver hepatocellular carcinoma; LUAD, Lung adenocarcinoma; OV, Ovarian serous cystadenocarcinoma; LUSC, Lung squamous cell carcinoma; PAAD, Pancreatic adenocarcinoma; PCPG, Pheochromocytoma and Paraganglioma; PRAD,Prostate adenocarcinoma; READ, Rectum adenocarcinoma; SARC, Sarcoma; SKCM, Skin Cutaneous Melanoma; STAD, Stomach adenocarcinoma; TGCT, Testicular Germ Cell Tumors; THCA, Thyroid carcinoma; THYM, Thymoma; UCEC, Uterine Corpus Endometrial Carcinoma; UCS, Uterine Carcinosarcoma. (**D**) Spearman correlation analysis revealing positive expression correlations between key identity-associated TFs and genes pivotal for both identity traits and cancer hallmarks in melanoma (top) and colorectal cancer (bottom), against their respective normal tissue counterparts, analyzed using GEPIA 2.0.

In order to investigate whether original cell identity is reinforced across cancer types, we selected both cancer types highly dependent on original cell identity, colorectal adenocarcinoma (CRC) and melanoma, to examine. CRC and melanoma respectively originate from intestinal stem cells and neural crest cells (*24–26*). We compared the expression of key identity-associated TFs between cancer cells and normal counterparts, including CTNNB1, TCF7L2, CDX2, and KLF5 for intestinal stem cells and SOX10, MITF, DUSP4, FOXD3, PAX3, and ETS1 for neural crest cells. Significantly, these TFs were consistently upregulated in their corresponding cancer cells (Fig. 4C), indicating a reinforced cell identities in both cancer types. Mechanistically, these reinforced identities are driven by genetic alterations (*3, 24, 26–31*).

We subsequently analyzed whether these reinforced identities endows cell with cancer hallmarks. Focusing on genes directly regulated by identity-associated TFs in intestinal stem cells and neural crest cells [29-31]. In colorectal adenocarcinoma cells, we observed a significant upregulation in these genes governing proliferation (c-myc, Cyclin D1, Cyclin E2, and VEGF) and maintaining an undifferentiated state (LGR5, Ascl2, Axin2, TCF7, CD44, SOX9, and Olfm4) compared to intestinal stem cells. Similarly, in melanoma cells, distinct gene sets involved in cell proliferation (MYC, BCL2, CDK2, MAPK, KIT, and CCND3), migration and invasion (SNAIL and SLUG), undifferentiation maintenance (ErbB3, TFAP2A, and MITF), and pigmentation (TYR, DCT, and TYRP1) (*27, 29, 32–35*) (*36, 37*), were expressed at higher levels relative to neural crest cells (fig. S8B). Importantly, these genes and their corresponding identity-associated TFs displayed positive expression correlations in colorectal adenocarcinoma and melanoma cells (Fig. 4D). These results indicated that reinforced intrinsic identities confer colorectal adenocarcinoma and melanoma cells with cancer features. Supporting this notion, knockdown of CTNNB1/CDX2/KLF5 in colorectal adenocarcinoma cells and Sox10/MITF in melanoma cells resulted in marked downregulation of these genes, significantly impairing cancer progression (*29, 31, 38–40*). In summary, reinforced cell identity is a pivotal driver of cancer hallmarks across cancer types.

### Identification of narciclasine as a disruptor of reinforced promonocyte identity in AML- _M5_MLL-AF9

Reinforced cell identity is a pivotal driver of cancer hallmarks across cancer types, implying disruption of reinforced cell identity as a novel strategy for cancer therapy. We continually focused on AML-M5^MLL-AF9^ to examine whether there are compounds disrupting reinforced promonocyte identity. Analyzing L1000 mRNA profiling data of AML-M5^MLL-AF9^ cells treated with compounds, we identified both candidate compounds, narciclasine and SCH-79797. Both compounds were able to significantly decrease the expression of key promonocyte master TFs (such as IRF8, SPI1, MYB, GFI1, or LMO2) (Fig. 5A) and their proliferation (Fig. 5B and fig. S9A). Furthermore, the effect of narciclasine and SCH79797 on inhibiting cell proliferation can be partially rescued by overexpressing GFI1 together with either SPI1 or IRF8 (Fig. 5C), indicating that both compounds inhibited AML proliferation through disruption of reinforced promonocyte identity. Narciclasine exhibited the most pronounced inhibitory effect on the expression of key promonocyte TFs. Consequently, we selected narciclasine for further investigations.

**Fig. 5.**
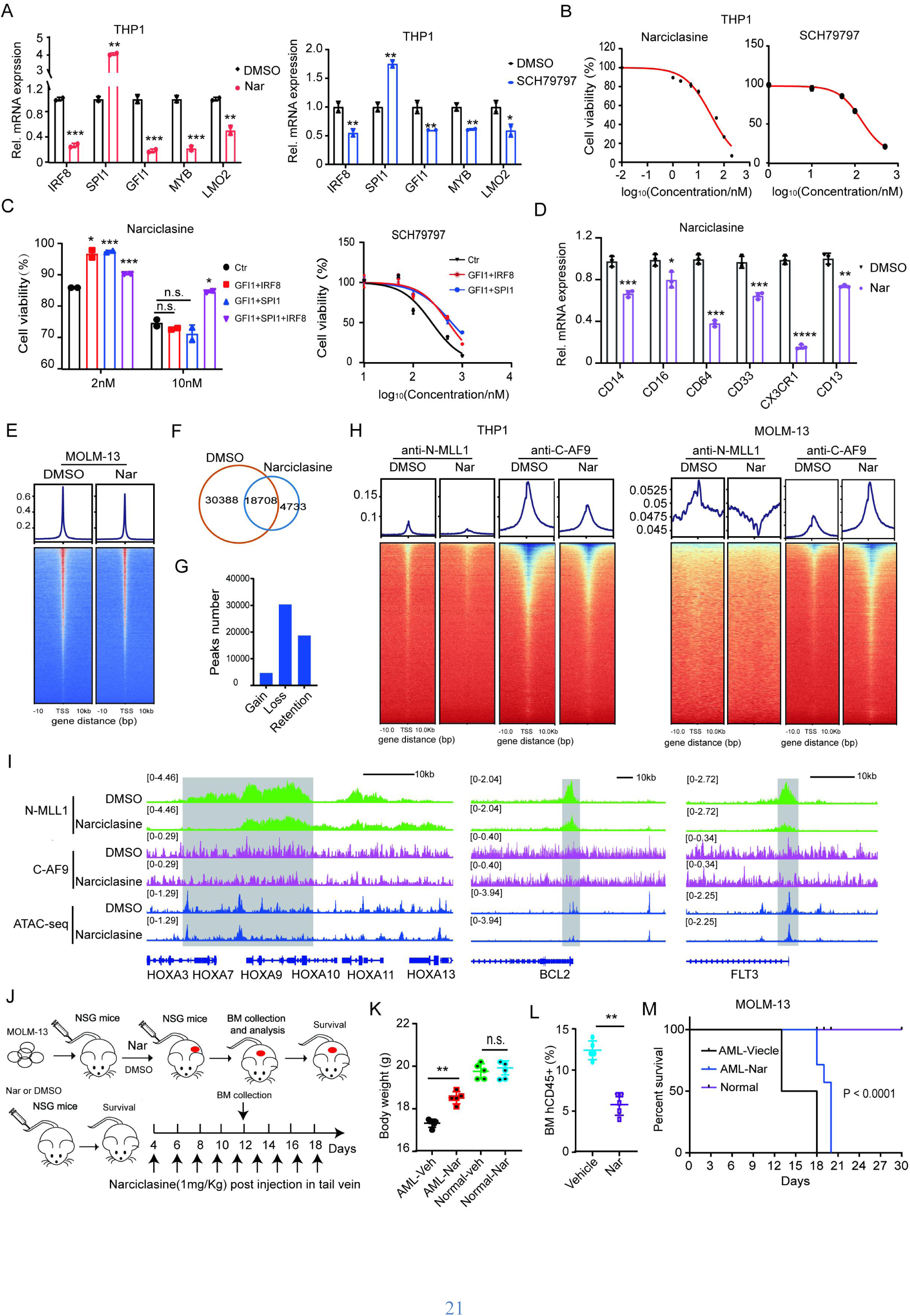
Identification of narciclasine as a disruptor of reinforced promonocyte identity in AML-M5^MLL-AF9^ cells (**A**) RT–qPCR results showing the expression levels of key promonocyte TFs in THP1 cells treated with DMSO, 50nM narciclasine or 1μM SCH-79797 treatment for 12h. (**B**) Dose response curves showing the inhibitory effects on THP1 cell numbers following treatment with indicated concentrations of narciclasine or SCH-79797 for 36h. (**C**) Bar graphs and dose response curves showcasing the impact on the survival of THP1 cells after 36h treatment with indicated concentration of narciclasine or SCH79797, along with overexpression of specified combinations of IRF8, SPI1 and GFI1. (**D**) RT-qPCR results displaying the expression levels of promonocyte cell surface markers in THP1 cells after treatment with 50 nM narciclasine (Nar) for 12h. (**E**) Metaplot (top) and density (bottom) plot illustrating ATAC-seq profiles in MOLM- 13 cells treated with DMSO or 50 nM narciclasine (Nar) for 12h. (**F**) Venn diagram showing the overlap of ATAC-seq peaks in MOLM-13 cells as described in (E). (**G**) Bar graphs revealing ATAC-seq peak changes in MOLM-13 cells as described in (E), categorized into peaks gained, lost, and unchanged. (**H**) Metaplot (top) and density (bottom) plot depicting ChIP-seq profiles for N-MLL1 and C-AF9 in THP1 (left) and MOLM-13 (right) cells treated with DMSO or 50 nM Narciclasine (Nar) for 12h. (**I**) Chip-seq and ATAC-seq gene tracks showing chromatin accessibility (top) and MLL-AF9 binding peaks (bottom) at selected loci in MOLM13 cells treated with DMSO or 50nM narciclasine (Nar) for 12h. (**J**) Schematic illustration of the workflow in evaluating in-vivo effects of narciclasine. (**K**) Analysis of body weights in NSG mice after transplantation with MOLM-13 cells and subsequent treatment with 100μL vehicle (AML-Veh) or 100μL Narciclasine (AML-Nar) every other day for 14 days. Control groups include NSG mice treated with vehicle (Normal-Veh) or Narciclasine (Normal-Nar) at the same dosage and regimen. (**L**) Flow cytometry analysis of the percentage of human CD45^+^ leukemia cells within the bone marrow of NSG mice, which were transplanted with MOLM-13 cells, and subsequently treated every other day for 14 days with 100μL vehicle (Vehicle) or 100μL Narciclasine (Nar) (n = 5 per group). (**M**) Kaplan-Meier survival curves for NSG mice as described in (m) (n = 6 per group). The p value was determined using a log-rank Mantel-Cox test.

Treatment with narciclasine led to significant reductions in the expression of most promonocyte surface markers (Fig. 5D; fig. S9, B and C) and loss of colonic capacity across both AML- M5^MLL-AF9^ cell lines (fig. S9D). ATAC-seq analyses revealed that narciclasine treatment resulted in a global reduction in chromatin accessibility, with a significant portion of the original 30388 open chromatin loci lost (61.8% loss), and only a few (4733) new open loci emerging in MOLM13 cells (Fig. 5, E to G). Subsequent ChIP-seq analysis showed that MLL-AF9’s chromatin binding was decreased following narciclasine treatment (Fig. 5H). Importantly, genes key for AML hallmarks (such as HOXA family, BCL2, FLT3, MYC, JMJD1C, MEIS1, GFI1, MEF2D and RUNX1) were situated at these loci and were significantly downregulated post- narciclasine treatment (Fig. 5I; fig. S9, E and F).

To access narciclasine’s *in-vivo* effects, we treated NSG mice transplanted with MOLM-13 cells with 1mg/kg narciclasine daily for 14 days. Consistent with previous reports (*23*), this regimen did not significantly affect the mice’s body weights and survival rate (Fig.5, J and K). Notably, narciclasine-treated mice exhibited a significant reduction in leukemia burden within the bone marrow, and a marked extension in survival time compared to untreated mice (Fig. 5, L and M). AML-M5^MLL-AF9^ cells isolated from the mice’s bone marrow a drastic downregulation of key promonocyte TFs as well as surface markers in narciclasine-treated mice relative to untreated controls (fig. S9G).

In conclusion, our findings demonstrated that narciclasine was able to effectively treat AML- M5^MLL-AF9^ by disrupting reinforced promonocyte identity.

## Discussion

How genetic mutations transform specific cell type remains a fundamental open question in cancer biology. Recent studies increasingly suggested that diver mutations initiate cancer by hijacking position identity, lineage factor PAX8, chromatin structure and signaling pathway (*3–6, 8*). However, it remains largely unclear how genetic alterations and specific cell type interact to influence cancer development. We investigated how interactions between genetic alterations and specific cell type established cancer state and then claim a paradigm that interactions between genetic alterations and specific cell type established a reinforced intrinsic cell identity conferring original cell with cancer hallmarks. According to this paradigm, we find compound to disrupt reinforced identity to treat cancer as a novel therapeutic strategy.

We discovered that MLL-AF9 enhances the expression of key promonocyte TFs, maintaining cell at promonocyte stage and then collaborates with these TFs to amplify the expression of TFs’ downstream genes, particularly key for AML hallmarks. Additionally, patients’ samples analysis revealed that key TFs of promonocyte identity are expressed at significantly higher levels in AML-M5^MLL-AF9^ cells compared to normal promonocytes. These results suggest that AML may emerge from reinforcement of intrinsic cell identity, challenging conventional view of AML development as arising from dedifferentiation of hematopoietic lineages (*20*). These results also pioneer to uncover the mechanism underlying tissue’s specificity of genetic alteration in in hematological cancer types, by demonstrating mechanism of MLL-AF9’ cell selectivity.

Extensive analysis revealed that melanoma and colorectal cancer cells also exhibit and depend on reinforced identities of neural crest progenitors and intestinal stem cells, respectively. These results suggest that the principle of reinforced intrinsic cell identity driving cancer hallmarks may be more general across cancer types and explain the comprehensive tissue specificity of genetic alterations.

We discovered that narciclasine effectively treat AML-M5^MLL-AF9^ by disrupting their reinforced promonocyte identity. This finding not just offers a promising therapeutic candidate for this AML subtype, but also illuminates a pathway towards developing novel cancer treatment by targeting reinforced intrinsic cell identities.

In summary, our study elucidates the previously an unrecognized tumor driver that reinforced cell identity established by interaction between genetic mutations and specific cell type confers cancer hallmarks, claim a new paradigm in cancer biology, and highlights the potential of targeting reinforced cell identity as an innovative anti-cancer drugs.

## Acknowledgments

We thank Zhimin Liu and Yuanyuan Yang for advice and discussions; Pengcheng Jiao and Jiaojiao Ji at FACS Core of Tsinghua University for flow cytometry assistance. We also thank Junke Zheng at the Key Laboratory of Cell Differentiation and Apoptosis of Chinese Ministry of Education, Shanghai Jiao Tong University School of Medicine for MLL-AF9 plasmid. Chu Yajing, State Key Laboratory of Experimental Hematology, Institute of Hematology, Chinese Academy of Medical Sciences for MLL-AF9::GFP plasmid.

## Author contributions

H.X.Y., P.Q.W. and P.C.T. contributed equally to this work and they have the right to be listed first in bibliographic documents. H.X.Y. conceived and performed experiments, analyzed and interpreted all the data, and wrote the paper. P.Q.W. constructed plasmids, analyzed tumor vulnerability and convervied experiments. P.C.T constructed plasmids, analyzed identity- associated compounds and performed qPCR experiments. Q.M.X. performed the luciferase experiments. H.W. and K.Z.H. cultured cells and performed transplant experiments. B.W.W. performed transplant experiments. D.W. ordered the reagents. T.H.M. and S.D. conceived experiments, supervised the research, interpreted results and wrote the paper. S.D. is the lead contact.

## Data and materials availability

This study did not generate new unique reagents. All raw sequencing and processed data has been deposited to the Gene Expression Omnibus (GEO) under accession GSE262254, GSE262255 and GSE262256. Additional information regarding materials and data analysis, as well as code required to reanalyze the data reported in this paper, may be obtained from the lead contact upon reasonable request, Seng Ding.

## Supplementary Materials

Materials and Methods Supplementary Text

Figs. S1 to S9 Tables S1 to S3

