## supplemental materials for "Reinforced identity confers cells with cancer hallmarks"

**The PDF file includes:**

Materials and Methods
Figs. S1 to S9
Tables S1 to S3

### Materials and Methods

#### Cell culture

THP-1 cells were cultured in RPMI-1640 supplemented with 10% Fetal Bovine Serum (GIBCO, 1009914) and 1% Penicillin/Streptomycin (P/S, GIBCO, 10378016). MOLM-13 cells were cultured in RPMI 1640 (GIBCO, C11875500BT) supplemented with 20% FBS and 1% P/S (GIBCO, 10378016). HEK293T cells were cultured in DMEM (GIBCO, C11995500BT) supplemented with 10% FBS (GIBCO, 1009914) and 1% P/S (GIBCO, 10378016). Cells were cultured at 37°C with 5% $CO_2$.

#### Mice

6-8 week old female NOD.Cg-Prkdc$^{scid}$Il2rg$^{tm1Wjl}$/SzJ (NSG) mice were purchased from the Beijing HUAFUKANG BIOSCIENCE Company. These mice were housed in group (maximal six mice per cage) under specific-pathogen-free conditions and maintained on a 12-hour reverse light-dark cycle with ad libitum access to food and water. All animal protocols were approved by the Institutional Animal Care and Use Committee at Tsinghua University.

#### sgRNA and plasmid cloning

All human sgRNAs were cloned by annealing the sense and antisense DNA oligonucleotides and ligating them into a BsmBI-digested lentiCRISPR v2 backbone (Addgene, #52961). Except for shRNAs of MLL-AF9 (LEP-shMLL-AF9.1039 and LEP-shMLL-AF9.643) purchased from Addgene, all other shRNAs are sourced from shRNA library at the Center of Biomedical Analysis, Tsinghua University. All sgRNA and shRNA sequences are listed in Table 2. Full length IRF8, SPI1, and GFI1 cDNA was amplified from a cDNA library prepared from THP1 cells, and individually cloned into a lentiviral expression vector, Lenti-EFS-T2A-Tdtomato, using the In-Fusion cloning system (HiFi DNA Assembly Master Mix, NEB #E2621). For MLL-AF9 construct, MLL-AF9 was PCR amplified from the pMIG-FLAG-MLL-AF9 plasmid (Addgene, #71443) and subcloned into a lentiviral expression vector, LentiEFS-T2A-EGFP, using the In-Fusion cloning system (HiFi DNA Assembly Master Mix, NEB #E2621).

#### Virus production and transduction

For lentivirus production, HEK293T cells were transfected with the plasmids encoding genes of interest, along with lentivirus packaging plasmids, psPAX2 and pMD2G. For transfecting cells in one well of a 6-well plates, 2μg plasmid encoding genes of interest, 2μg pPAX2, and 1μg pMD2G were mixed with Lipofectamine 3000 (Invitrogen, L3000008) in 250μL OPTI-MEM

(GIBCO, 31985-070). For cells in a 10-cm plate, the transfection mixture contained 10μg plasmid encoding genes of interest, 10μg pPAX2, 5μg pMD2G, and Vigofect (Vigororous, T001) in 800μL OPTI-MEM. Transfected HEK293T were incubated for 6 hours before media was removed and replenished with fresh media. Lentivirus was harvested at 24 and 48 hours post-transfection, with the collected virus pooled for subsequent use. For retrovirus production, PlatA cells were transfected with the plasmid of interest for 10cm plates, transfection mixture comprised 15μg plasmid encoding genes of interest and Lipofectamine 3000 (Thermo Fisher Scientific, L3000) in 800μL OPTI-MEM (GIBCO, 31985-070). Virus was collected at 24 and 48 hours post-transfection and pooled together. For viral transduction, filtered virus-containing supernatant were applied to different cell lines. Medium was replenished at 24 hours post-infection, and supplemented with 4μg/mL (in THP1 cells) or 1.5ug/mL (in MOLM-13 cells) puromycin (Solarbio, P8230) for subsequent selection.

**Cell viability assay**

To evaluate the effect of knocking out/down IRF8, SPI1 or GFI1 on the cell growth, cells both with and without genetic manipulations, were plated in opaque-walled 96-well plates at a density of 800 cells per well. These cells were cultured for periods of 1, 2, 3, 5, and 7 days. Subsequently, cell viability was measured with a Microplate Reader (PerkinElmer EnSpire) using Cell Titer Glo Luminescent Cell Viability Assay kit (Beyotime Biotechnology, C0065L) following the manufacturer's instructions.

**Cell cycle analysis**

THP1 and MOLM-13 cells both with and without knockdown of IRF8, SPI1, or GFI1, were counted. $2 \times 10^5$ cells from each group were centrifuged at 200g for 5 min. The cell pallets were then washed with DPBS (Thermo Fisher Scientific, C14190500BT). After resuspension of the pellets in 300μL DPBS, 700μL cold ethanol were added slowly with vortexing, ensuring thorough fixation and minimizing cell clumping. Following fixation for overnight at 4°C, cells were wash twice with DPBS, and then centrifuged at 850 g for 4 min. The supernatants were carefully removed, and the cell pellets were resuspended in 100 μL of DPBS. For DNA staining, 50μL of 100 μg/mL RNase A (Thermo Fisher Scientific, EN0531) and 200 μl 50 μg/mL propidium iodide (PI, MCE, HY-D0815). The cell-cycle distribution in each sample was subsequently analyzed on a BD FACS Aria III cell sorter. The resulting data were analyzed using FlowJo10 software.

**Colony formation assay**

To assess the colony formation capacity of THP1 and MOLM-13 cells after specific genetic manipulations, cells were collected and washed with IMDM (Thermo Fisher Scientific, C12440500BT) supplemented with 2% FBS and 1% P/S. Then cells were seeded into xx plates using MethoCult$^{TM}$ H3434 Classic medium (Stem Cell Technologies, #04434) at density of 20,000 cells per well. Following 10 days of incubation, colonies formed from each sample were counted using Hemocytometer (Fisher, 14921490 3100) according to the manufacturer's instructions.

**Screening of small-molecule cell identity disruptors**

Compounds that trigger the death of AML cells were identified using the DepMap and L1000 databases. Transcriptional analysis was conducted on AML cells treated with these compounds for either 6 or 8 hours, and the outcomes were presented in a heat map format. Small-molecule cell identity disruptors selected from an initial DepMap and L1000 library-based screening were further validated by measuring their inhibitory effects on the expression of promonocyte identity genes. THP1 or MOLM13 cells were plated onto 96 well plates, treated with the compounds at from 1nM to 10μM for 36hrs and DMSO as a basal level control. Cell viability was monitored using Cell Titer-Glo Luminescent Cell Viability Assay (Beyotime Biotechnology, C0065L). THP1 or MOLM13 cells were plated onto 6 well plates treated 50nM narciclasine, 100nM SH 79797 and 500nM Piperlongumine for 12h and then tested for the expression of identity genes by qPCR.

**Apoptosis analysis**

Apoptosis in THP1 and MOLM-13 cells, with or without knockdown of IRF8, SPI1, or GFI1, were quantified via flow cytometry. Briefly, counted $2 \times 10^5$ cells from each condition were centrifuged at 200 g for 5 min. The cell pallets were resuspended in 100μL of FACS buffer (DPBS + 5%FBS) and stained with 5μL Annexin V, followed by 5μL PI, both from Apoptosis kit (Absin, abs50001-50T), for 15min and 5 min respectively. After three washes with DPBS and subsequent centrifugation at 200 g for 5 min, the cells were resuspended in 300μL of FACS buffer. The percentage of PI and Annexin V-positive cells were determined using a BD Aria III FACS cell sorter, with data analyzed via FlowJo10 software.

**AML-M5$^{MLL\text{-}AF9}$ xenograft mouse models and narciclasine treatment**

The NSG mice, females aged 6-8 weeks, were used to establish AML-M5$^{MLL-AF9}$ xenograft model by intravenous injection of $2 \times 10^6$ MOLM-13 cells per mouse. 12 days post injection, mice exhibiting severe disease phenotypes, characterized by over 60% of peripheral blood cells being hCD45$^+$, received a tail vain administration of 1mg/kg narciclasine every other day for 2 weeks. To assess the impact of narciclasine, peripheral blood was collected via retro-orbital bleeding and bone marrow was harvested by flushing femurs and tibias with PBS following sacrifice. Samples were treated with red blood cell lysis buffer (Roche, 11814389001) on ice for 5 min as per manufacturer's instruction and centrifuged at 200g for 5 min. The Pellets were then resuspended in 300 μL of PBS and filtered through 70μm sterile strainers (Corning, 352350). Cells were stained with anti-human CD45 (1:100, BioLegend, 304007) and anti-mouse CD45 (1:100, BioLegend, 103116) antibodies. The percentage of CD45-positive cells were determined using a BD AriaFACS cell sorter, and the data analyzed by FlowJo10 software.

**Construction of reporter system and luciferase assay**

To investigate the promoter activity of genes bound by MLL-AF9, a series of 1.5-2 kb DNA sequences encompassing 4.5-6 kb promoter regions upstream of IRF8, SPI1, GFI1, and CEBPA (listed in Table 3), was cloned from genomic DNA isolated from THP1 cells. Each DNA sequence was individually cloned into a PGL4.23 plasmid using ClonExpress® Ultra One Step Cloning Kit (Vazyme Biotech, C115-02). For luciferase assay, HEK293T cells were co-transfected with the constructed reporter plasmids, along with plasmids encoding MLL-AF9 and Renilla luciferase. Firefly and Renilla luciferase activities were measured 24 hours post-transfection using the Dual-Luciferase Reporter Assay System (Beyotime Biotechnology, RG027) on a Microplate Reader (PerkinElmer EnSpire, 2300) following the manufacturer's instructions.

**Western blot analysis**

Proteins were extracted from THP1 and MOLM-13 cells using RIPA buffer (Beyotime Biotechnology, P0013B) supplemented with protease and phosphatase inhibitors (Beyotime Biotechnology, P1045). Blots were incubated with primary antibodies against β-actin (1:1000; Santa Cruz Biotechnology, sc-47778), N-MLL1 (1:3000; Bethyl Laboratories, A300-087A), AF9 (1:2000; Thermo Fisher Scientific, PA5-27797), HOXA7 (1:1000; Santa Cruz Biotechnology, sc-81290), FLT3 (1:1000; Santa Cruz Biotechnology, sc-19635), c-MYC (1:1000; Santa Cruz Biotechnology, sc-40), BCL2 (1:1000; Santa Cruz Biotechnology, sc-7382), IRF8 (1:1000; Invitrogen, PA5-85224), SPI1 (1:1000; Cell Signaling Technology, #2266s), GFI1 (1:1000;

Santa Cruz Biotechnology, sc-373960), CEBPA (1:1000; Santa Cruz Biotechnology, sc-166258). For detection, HRP-linked goat anti-mouse (1:5000; Santa Cruz Biotechnology, sc-516102), and mouse anti-rabbit (1:5000; Santa Cruz Biotechnology, sc-2357) secondary antibodies were applied. Blots were developed using ECL (GE Healthcare) as per the manufacturer's instructions.

**Immunoprecipitation**

For immunoprecipitation, cells cultured in 10 cm dishes to approximately 90% confluency were washed once with cold DPBS, and harvested by trypsinization. Following centrifugation at 1400rpm for 5 min, the cell pellets were lysed using 5-10 times their volumes of NP40 lysis buffer (Beyotime Biotechnology, P0013F), and incubated on the ice for 30 min. Post-lysis, the lysates were centrifuged at 12, 000g for 15 min, and the supernatants were transferred to new tubes for protein concentration measurement using Pierce™ BCA Protein Assay Kits (Thermo Fisher Scientific, 23227), as per manufacturer's instructions. Samples were then adjusted to a protein concentration of 2mg/mL in a total volume of 500μL with NP40 lysis buffer. From these, 50μL of each sample was set aside as input, and the remaining samples were mixed with specific antibodies and incubated at 4°C for 8 hours. Subsequently, 20 μL of protein A/G magnetic beads (Merck Millipore, LSKMAGAG10) were added, and the mixture was incubated at 4°C for an additional 4hr. Beads were washed for 3 times with NP40 lysis buffer, resuspended in 1 × SDS loading buffer, boiled, and subjected to Western blot analysis. The antibodies used in immunoprecipitation included anti-N-MLL1 (1:200; Bethyl Laboratories, A300-087A), anti-AF9 (1:200; Bethyl Laboratories, A301-567A), anti-IRF8 (1:50, Abcam, ab98166), anti-SPI1 (1:100; Cell Signaling Technology, #2266s), anti-GFI1 (1:50; Santa Cruz, sc-373960) and IgG isotype control (1:200; Abcam, ab171870).

**RT-qPCR analysis**

RNA was extracted from cell samples using the Axyprep multisource total RNA kit (Axygen, AP-MN-MS-RNA-250), following the manufacturer's instructions. From each sample, 1μg of total RNA was reverse transcribed into cDNA using the EasyScript® All-in-One First-Strand cDNA Synthesis SuperMix (TransGen biotech, AE341-02), according to manufacturer's instructions. Subsequent RT-qPCR analyses were performed using PerfectStart® Green qPCR SuperMix (TransGen biotech, AQ601-01-V2) on a Bio-Rad CFX96 system. The Ct values obtained were normalized against Actin to quantify relative gene expression levels. The sequences of all RT primers utilized were listed in Table 4. For treatment involving small molecules, 1 × $10^6$ MOLM-13 or THP1 cells were treated with 50nM narciclasine, 200nM SCH 79797 or 500nM piperlongumine for 36h prior to RNA extraction. Cells treated with 0.05% DMSO served as the control group.

**RNA-seq**

For RNA-seq analysis of MOLM-13 and THP1 cells with or without knockdown of IRF8, SPI1, GFI1 or MLL-AF9, cells were collected on day 4 post-infection of viral shRNA plasmids. Total RNA was extracted from 1.5 × $10^7$ cells of each cell sample using the Axyprep multisource total RNA kit (Axygen, AP-MN-MS-RNA-250). The RNA-seq libraries were constructed using the using NEBNext® UltraTM RNA Library Prep Kit for Illumina® (#E7530L, NEB) according to manufacturer's instructions. Briefly, 1μ of total RNA underwent poly-A selection and fragmentation using the NEBNextPoly(A) mRNA Magnetic Isolation Module (#7490, NEB), followed by first and second-strand cDNA synthesis. The libraries were then 150bp end-repaired, adaptor-ligated with the Illumina adaptors, and PCR amplified with 8 cycles. Sequencing was performed on Illumina Novaseq 6000 platform.

**ChIP-seq**

ChIP-seq analysis was performed on THP1 and MOLM-13 cells, with or without specific genetic manipulations, using Chromatin Immunoprecipitation kit (Millipore, 17-371) as per the manufacturer's instructions. Briefly, a total of 2 x $10^7$ cells from each experimental condition were harvest 5 days post-knockdown/knockout procedure. The cells were fixed with 1% formaldehyde for 10 min at room temperature to achieve cross-linking, and quenched with 0.125 M glycine for an additional 10 min. Post-fixation, cells were washed with DPBS and lysed in 130μL of SDS Lysis Buffer (Beyotime Biotechnology, P0013G) supplemented with 1 X Protease Inhibitor (Beyotime Biotechnology, P1045). The lysates were subjected to sonication using a Covaris S220 Focused-ultrasonicator. The sonication conditions were set to 150W Peak Incident Power, 10% Duty Factor, 200 Cycles per Burst, for 120s. This is followed by centrifugation at 14,000 g for 15 min at 4°C. Chromatin containing supernatants (100μL) were transferred to new tubes, and mixed with 900 μL of Dilution Buffer (20mM Tris, pH 8.0, 2mM EDTA, 150mM NaCI, 1% Triton X-100, and 0.01% SDS) supplemented with 1 X Protease Inhibitor (Beyotime Biotechnology, P1045). A 10 μL aliquot from each sample was reserved as input. The remaining supernatant were incubated overnight at 4°C with rotation, using 2μL of anti-N-MLL1 (Bethyl Laboratories, A300-087A), 3uL of anti-AF9 (Thermo Fisher Scientific,

PA5-27797), 10 μL of anti-IRF8 (Cell Signaling Technology, 83413T), 10μL of anti-SPI1 (Cell Signaling Technology, #2266s), or 10μL of anti-GFI1 (Santa Cruz Biotechnology, sc-373960) antibodies. The mixture was then incubated with 40μL protein A/G magnetic beads (Merck Millipore, LSKMAGAG10) for 4h at 4°C. After incubation, beads were washed sequentially with Low Salt, High Salt, LiCl Immune Complex Wash Buffers (Millipore, 17-371), and twice with TE buffer (pH 8.0). Chromatin was eluted in 200μL elution buffer (1μL 20% SDS, 20μL 1M $NaHCO_3$ and 170 μL sterile-distilled water) with shaking at 600 rpm at 65°C for 15min. The eluted chromatin was reverse-crosslinked with 0.25M NaCI overnight at 65°C, treated with 1μL of RNase A (Thermo Fisher Scientific, EN0531), 1μL Proteinase K (Cell Signaling Technology, 10012S) together with 4μL 0.5 M EDTA and 8μL 1M Tris-HCl at 37°C for 30min. DNA was then purified using QIAquick PCR purification kit (QIAGEN, 28104), following the manufacturer's instructions. ChIP-seq libraries were prepared using NEBNextUltra II DNA Library Prep Kit for Illumina (NEB, #E7103), according to the manufacturer's instructions. The libraries were then 150bp end-repaired, adaptor-ligated with the Illumina adaptors, and PCR amplified with 8 cycles. Sequencing was performed on Illumina Novaseq 6000 platform.

**ATAC-seq**

MOLM-13 cells, treated with 50nM narciclasine or 0.05% DMSO for 12h and THP1 and MOLM-13 cells subjected to knockout or knockdown of IRF8, SPI1, or GFI1 for 3 days, were harvested for ATAC-seq analysis. The ATAC-seq library preparation was performed. In brief, after cells were washed by 1mL DPBS, 50,000 cells were centrifuged at 500g for 5 min in a 4 °C pre-chilled fixed-angle centrifuge and washed by ice-cold PBS. After centrifugation, supernatant was removed with two pipetting steps to avoid the cell pellet. Cell pellets were then resuspended in 50 μl of ATAC-seq RSB containing 0.1% IGEPAL-630, 0.1% Tween-20, and 0.01% digitonin and incubated on ice for 10 min. After lysis, 1 ml of ATAC-seq RSB containing 0.1% Tween-20 (without IGEPAL-630 or digitonin) was added, and the tubes were inverted for six times to mix. Nuclei were then centrifuged for 10 min at 500g in a 4 °C pre-chilled fixed-angle centrifuge. Supernatant was removed and nuclei were then incubated with the Tn5 transposome and tagmentation buffer at 37 °C for 30 min (Novoprotein, N248). After the tagmentation, the stop buffer was added directly into the reaction to end the tagmentation. Reactions were cleaned up with 2.2x VAHTS DNA Clean Beads (Vazyme, N411). PCR was performed to amplify the library for 8 cycles by KAPA HiFi PCR Kit (Kapa Biosystems, kk2102) using the following

PCR conditions: 72 °C for 5 min; 98 °C for 3mins; and thermocycling at 98 °C for 20 s, 63 °C for 30 s and 72 °C for 3 min. After the PCR reaction, libraries were purified with the 1.8x VAHTS DNA Clean Beads.

**Analysis of RNA-seq, ChIP-seq and ATAC-seq data**

For RNA-Seq data analysis, sequencing reads were aligned to the human reference genome (GRCh38/hg38) using STAR Aligner (V2.7.3a), employing default parameters. Following alignment, raw read counts were generated using featureCounts (subread-2.0.3) from the Subread package. RPM (reads per million)-normalized bigwig files were created using bedGraphToBigWig (UCSC), allowing for visualization in the Integrative Genomics Viewer (IGV) (version 2.14.1). Mapped reads were analyzed with DESeq2 (version 1.14.1) to identify differentially expressed genes.

ChIP-seq data analysis involved aligning sequencing reads to human reference genome. Reads from the GEO database (GSE123872 and GSE79899) were aligned to the human genome GRCh37/hg19 using Bowtie2 (version 2.3.5). Similarly, sequencing reads from this study, derived from THP1 and MOLM-13 cells with or without specific genetic manipulations, were aligned to human genome GRCh38/hg38 using Bowtie2 (version 2.3.5). Presumed PCR duplicates were removed using Picard tools (version 1.96) by employing the MarkDuplicates command. ChIP-Seq signal peaks were identified using MACS (version 2.1), applying the NarrowPeak setting with a q value cutoff of 0.05. Specifically for N-MLL1 ChIP-Seq, signal peaks were called on using MACS v2.1 with both NarrowPeak and broadPeak settings utilizing a p value cutoff of 1e-8 and a broad p value cutoff 1e-4, respectively. Genes proximal to identified peaks were annotated based on the GRCh37/hg19 or GRCh38/hg38 genome assembly using ChIPpeakAnno package from Bioconductor. Venn diagram to compare peak overlaps were plotted using the same ChIPpeakAnno package. Heatmaps and metaplots of binding signals were generated with the plotHeatmap function from deepTools suite. CPM (counts per million)-normalized bigwig files were created using both bedGraphToBigWig (UCSC) and bamCoverage (deepTools). These bigwig files facilitated the visualization of binding signals in IGV 2.14.1.

For ATAC-seq data analysis, sequencing reads were aligned to the human reference genome (GRCh38/hg38) using Bowtie2 (version 2.3.5). Picard tools (version 1.96) was employed to remove presumed PCR duplicates with the MarkDuplicates command. ATAC-seq signal peaks were called using MACS (version 2.1) with the following command: macs2 callpeak –t -n sample --shift -100 --extsize 200 --nomodel -g hs. The venn diagrams comparing peaks were plotted using Bioconductor package ChIPpeakAnno. Heatmaps and metaplots were generated with the plotHeatmap function from the deepTools suite. CPM-normalized bigwig files were created using both bedGraphToBigWig (UCSC) and bamCoverage (deepTools). These bigwig files were then used to visualize binding signals in IGV 2.14.1.

**Statistical analysis.**

The experiments in this study were performed with at least three biological replicates unless specified. All statistical analyses were performed with Graphpad prism 8.0 software or R Bioconductor. General statistical analyses were performed using a two-tailed Student's t-test for data with normal distribution and equal variances. For datasets not meeting these criteria, the Wilcoxon test was applied. Both tests were utilized to make inferences at a 95% confidence interval unless specified otherwise. The tests used and the P-values are listed on figures and figure legends. Details of statistical tests are outlined within figures and figure legends. *$P < 0.05$, ** $P < 0.01$, ***$P < 0.001$, ****$P < 0.0001$.

**Data availability.**

The raw sequencing datasets generated in this study are available in the Gene Expression Omnibus (GEO) database, under the accession number (GSE262254, GSE262255 and GSE262256). All other data supporting the findings of this study are available from the corresponding author upon reasonable request.

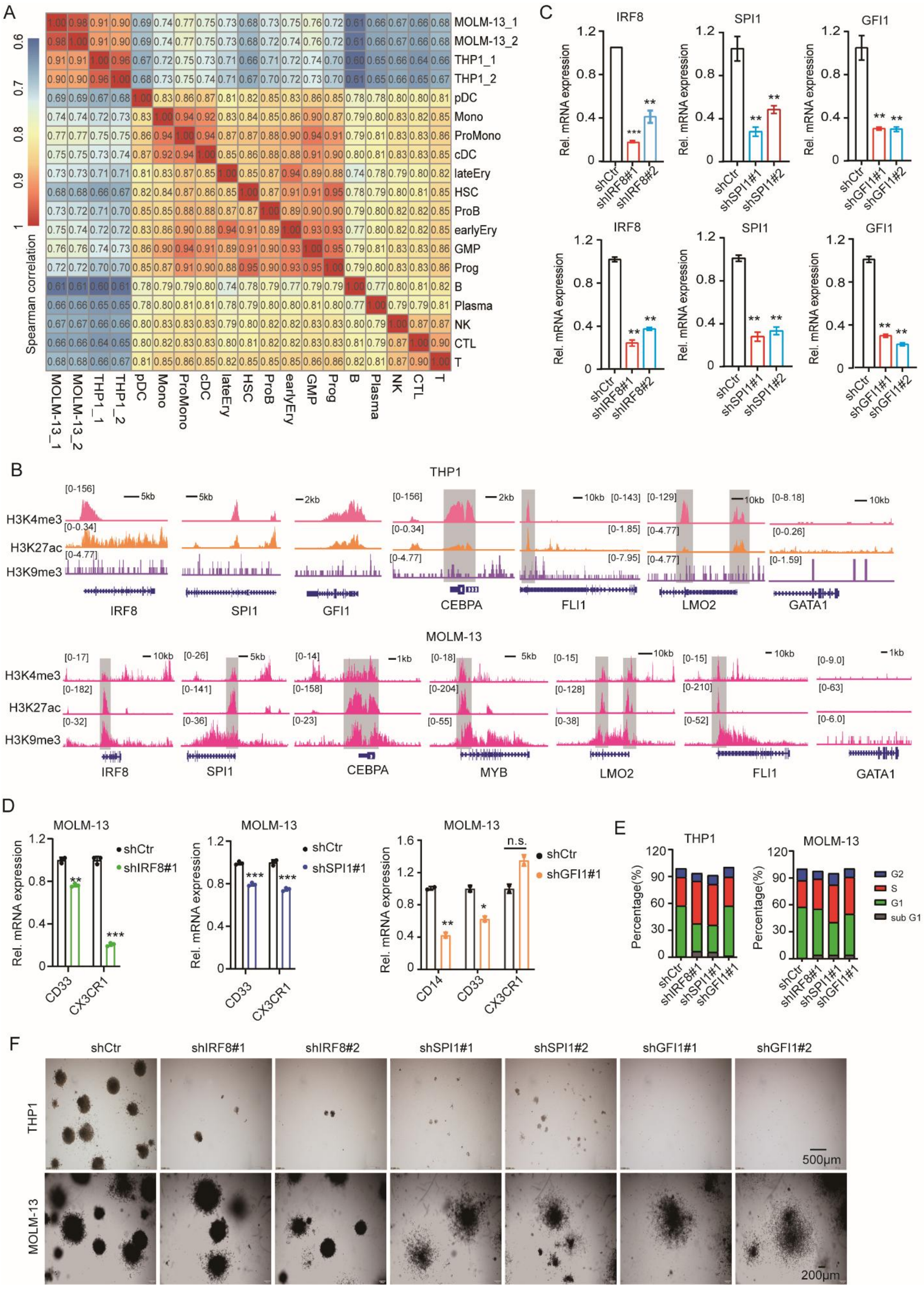
A
MOLM-13_1
MOLM-13_2
THP1_1
THP1_2
pDC
Mono
ProMono
cDC
lateEry
HSC
ProB
earlyEry
GMP
Prog
B
Plasma
NK
CTL
T
Spearman correlation
C
IRF8
SPI1
GFI1
Rel. mRNA expression
shCtr
shIRF8#1
shIRF8#2
shSPI1#1
shSPI1#2
shGFI1#1
shGFI1#2
B
THP1
H3K4me3
H3K27ac
H3K9me3
IRF8
SPI1
GFI1
CEBPA
FLI1
LMO2
GATA1
MOLM-13
MYB
D
MOLM-13
shCtr
shIRF8#1
shSPI1#1
shGFI1#1
Rel. mRNA expression
CD33
CX3CR1
CD14
n.s.
E
THP1
MOLM-13
Percentage(%)
G2
S
G1
sub G1
F
shCtr
shIRF8#1
shIRF8#2
shSPI1#1
shSPI1#2
shGFI1#1
shGFI1#2
THP1
MOLM-13
500μm
200μm

**Fig. S1 AML-M5$^{MLL-AF9}$ cells present and depend on promonocyte identity**

(**A**) Correlation analysis between THP1 and MOLM-13 cells and various indicated normal hematological cell lineages. pDC, Plasmacytoid dendritic cells; Mono, Monocyte; Promono, Promonocyte; cDC, Classical Dendritic Cells; lateEry, late Erythrocytes; HSC, Hematopoietic stem cells; earlyEry, early Erythrocytes; GMP, granulocyte/monocyte progenitor; NK, Natural killer cells; CTL, Cytotoxic T lymphocytes. (**B**) ChIP-seq gene tracks showing the distribution of H3K27ac, H3K4me3, and H3K27me2 marks at indicated gene loci in THP1 (top) and MOLM-13 (bottom) cells, utilizing data from GSE79899 and GSE157636, respectively. (**C**) RT-qPCR results validating specified shRNA-mediated knockdown of IRF8, SPI1, and GFI1 in THP1 (top) and MOLM-13 (bottom) cells, with scramble shRNA (shCtr) serving as a control. (**D**) RT-qPCR analysis indicating changes in the expression of indicated promonocyte surface markers in MOLM-13 cells upon knockdown of IRF8, SPI1, or GFI1 using specified shRNA. (**E**) Quantitative analysis for FACS showing the cell cycle distribution in THP1 and MOLM-13 cells transduced with scramble control shRNA (shCtr) or shRNAs targeting IFR8, SPI1 or GFI1 for 72h. (**F**) Bright-flied images displaying the clones formed in methylcellulose culture by THP1 (top) and MOLM-13 (bottom) cells transduced with scramble control shRNA (shCtr) or specified shRNAs against IFR8, SPI1 or GFI1 for 7 days.

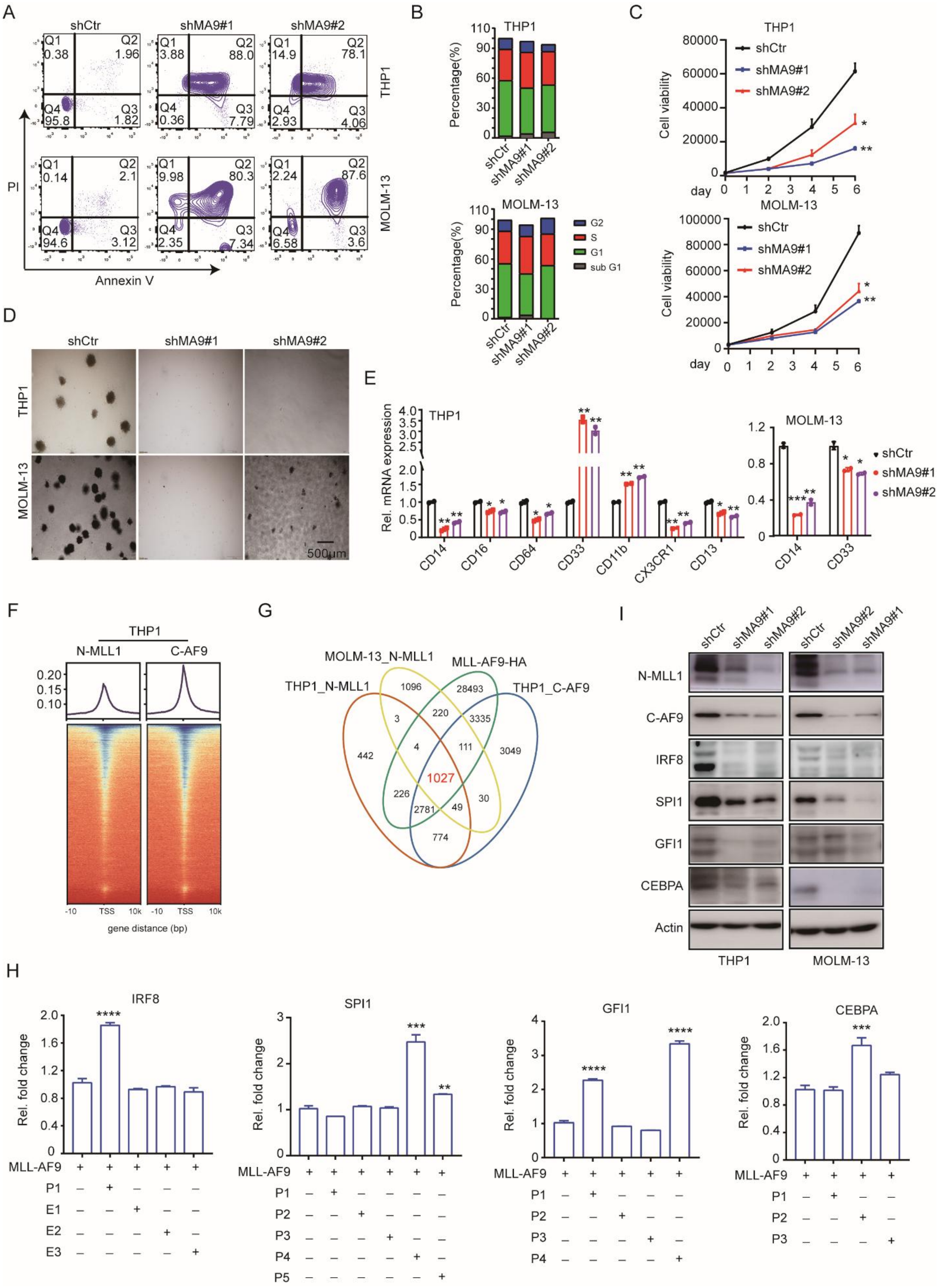
A
shCtr
shMA9#1
shMA9#2
Q1 0.38
Q2 1.96
Q4 95.8
Q3 1.82
Q1 3.88
Q2 88.0
Q4 0.36
Q3 7.79
Q1 14.9
Q2 78.1
Q4 2.93
Q3 4.06
THP1
Q1 0.14
Q2 2.1
Q4 94.6
Q3 3.12
Q1 9.98
Q2 80.3
Q4 2.35
Q3 7.34
Q1 2.24
Q2 87.6
Q4 6.58
Q3 3.6
MOLM-13
PI
Annexin V
B
THP1
Percentage(%)
shCtr
shMA9#1
shMA9#2
MOLM-13
G2
S
G1
sub G1
C
THP1
shCtr
shMA9#1
shMA9#2
Cell viability
day
MOLM-13
D
shCtr
shMA9#1
shMA9#2
THP1
MOLM-13
500μm
E
THP1
Rel. mRNA expression
CD14
CD16
CD64
CD33
CD11b
CX3CR1
CD13
MOLM-13
shCtr
shMA9#1
shMA9#2
F
THP1
N-MLL1
C-AF9
TSS
gene distance (bp)
G
MOLM-13_N-MLL1
MLL-AF9-HA
THP1_N-MLL1
THP1_C-AF9
1096
28493
3
220
3335
442
4
111
3049
1027
226
2781
49
30
774
I
shCtr
shMA9#1
shMA9#2
shCtr
shMA9#2
shMA9#1
N-MLL1
C-AF9
IRF8
SPI1
GFI1
CEBPA
Actin
THP1
MOLM-13
H
IRF8
Rel. fold change
MLL-AF9
P1
E1
E2
E3
SPI1
P2
P3
P4
P5
GFI1
CEBPA

**Fig. S2. MLL-AF9 maintains promonocyte identity by enhancing key TFs expression in AML-M5$^{MLL\text{-}AF9}$ cells**

(**A**) Apoptosis in THP1 and MOLM-13 cells transduced with scramble (shCtr) and two different MLL-AF9 (shMA9#1 and shMA9#2) shRNAs for 72h. (**B**) Quantitative analysis for FACS displaying the cell cycle distribution in THP1 and MOLM-13 cells after transduction with control scramble (shCtr) or ML-AF9 specific (shMA9#1 and shMA9#2) shRNAs for 72h. (**C**) Cell proliferation curves depicting the number of THP1 and MOLM-13cells at specified days post-transduction with scramble (shCtr) or MLL-AF9-targeting (shMA9#1and shMA9#2) shRNAs. (**D**) Bright-field images showing the clones formed in the methylcellulose culture by THP1 (top) and MOLM-13 (bottom) cells transduced with control scramble (shCtr) or ML-AF9-targeting (shMA9#1 and shMA9#2) shRNAs for 7 days. (**E**) RT-qPCR analysis of the expression levels of indicated promonocyte surface markers in THP1 and MOLM-13 cells following transduction with scramble (shCtr) and MLL-AF9 (shMA9#1 and shMA9#2) shRNAs for 72h. (**F**) Metaplot (top) and density plot (bottom) illustrating ChIP-seq profiles for N-MLL1 and C-AF9 in THP1 cells. (**G**) Venn diagram showing the overlap of binding peaks for N-MLL1, C-AF9, and MLL-AF9-HA in THP1 and MOLM-13 cells, using data from GSE173599. The number of overlapped MLL-AF9 peaks between THP1 and MOLM-13 cells was highlighted in red. (**H**) Reporter assay results illustrating the transcriptional activities of the sequences from MLL-AF9-bound chromatin marked by gray boxes in (fig. S3A). Data were derived from 293T cells with MLL-AF9 overexpression for 24h. (**I**) Western blot analyses showing the expression levels of key promonocyte TFs in THP1 and MOLM-13 cells following transduction with control scramble (shCtr) or MLL-AF9 specific (shMA9#1 and shMA9#2) shRNAs for 72h.

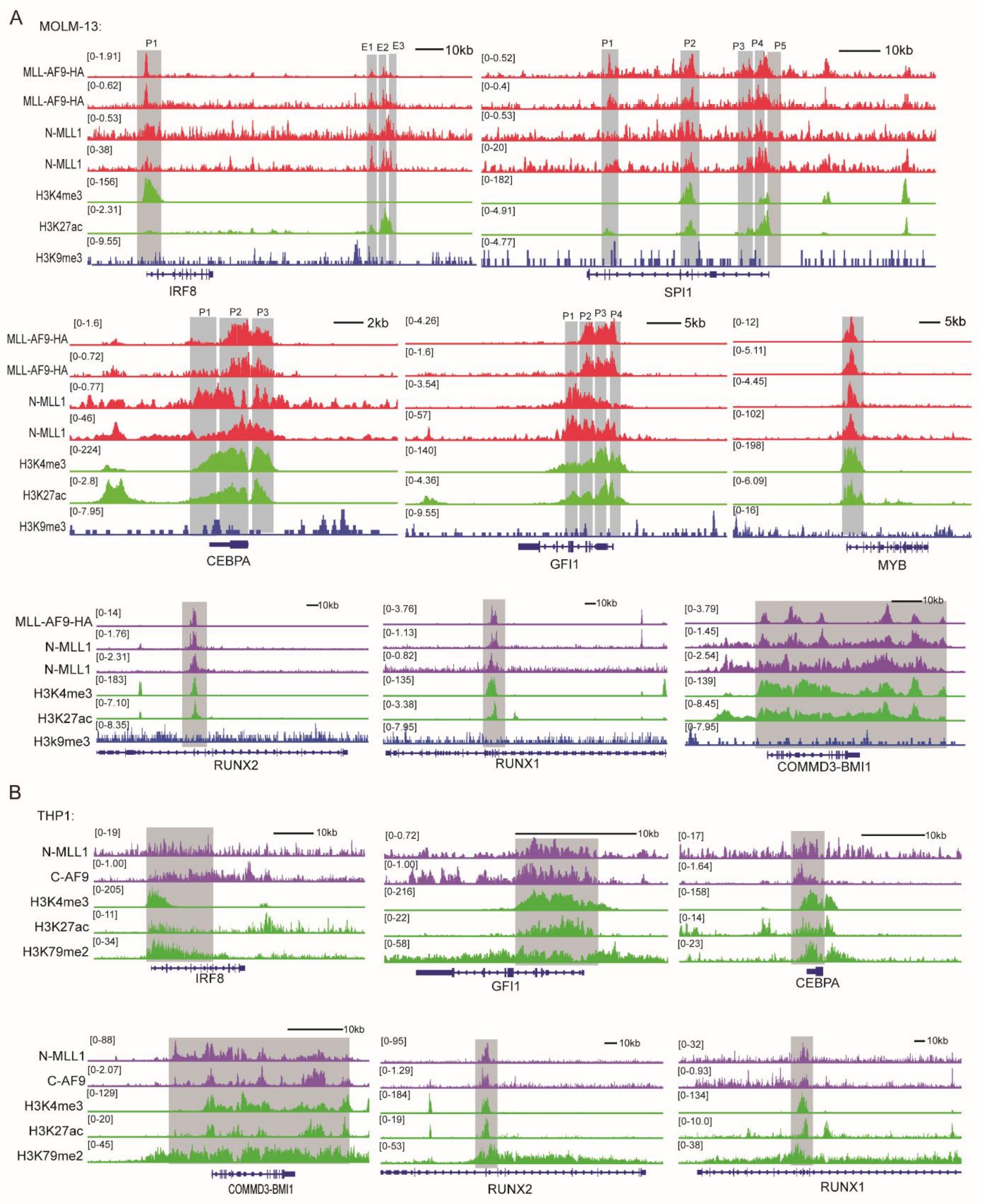
A
MOLM-13:
P1
E1 E2 E3
10kb
MLL-AF9-HA
MLL-AF9-HA
N-MLL1
N-MLL1
H3K4me3
H3K27ac
H3K9me3
IRF8
P1
P2
P3 P4 P5
SPI1
P1 P2 P3
2kb
CEBPA
P1 P2 P3 P4
5kb
GFI1
MYB
RUNX2
RUNX1
COMMD3-BMI1
B
THP1:
N-MLL1
C-AF9
H3K4me3
H3K27ac
H3K79me2
IRF8
GFI1
CEBPA
COMMD3-BMI1
RUNX2
RUNX1

**Fig. S3 MLL-AF9 binds to promoters of key promonocyte TFs in AML-M5$^{MLL\text{-}AF9}$ cells**

(**A**) ChIP-seq gene tracks displaying the chromatin binding of MLL-AF9-HA, N-MLL1 (from GSE173599), N-MLL1 replicate, and epigenetic marks H3K4me3, H3K27ac, and H3K9me3 at indicated gene loci in MOLM-13 cells. (**B**) ChIP-seq gene tracks revealing the chromatin binding of N-MLL1 and C-AF9, and epigenetic marks H3K4me3, H3K27ac and H3K79me2 at indicated gene loci in THP1 cells, using data from GSE79899.

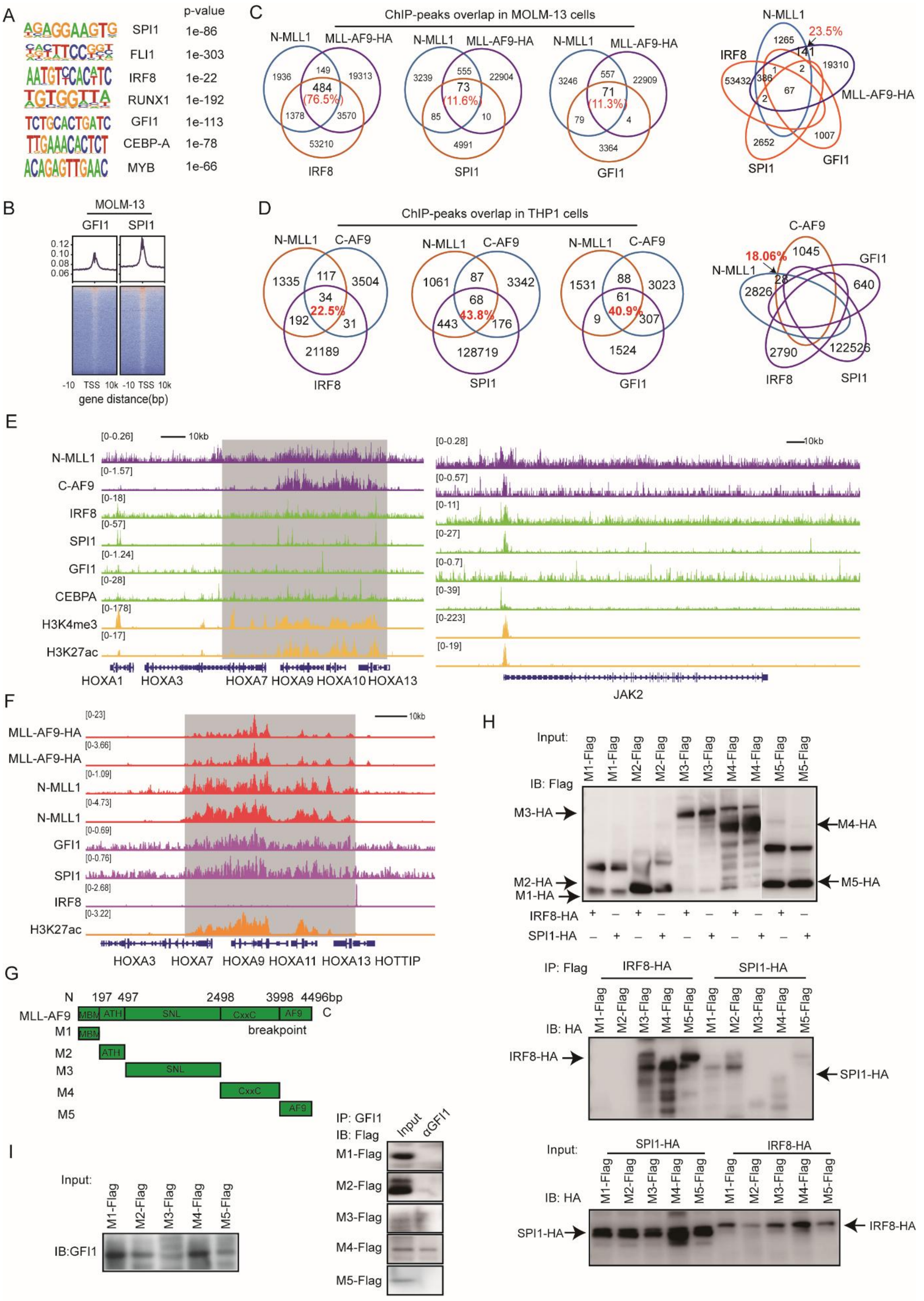
A
p-value
SPI1 1e-86
FLI1 1e-303
IRF8 1e-22
RUNX1 1e-192
GFI1 1e-113
CEBP-A 1e-78
MYB 1e-66
B
MOLM-13
GFI1 SPI1
gene distance(bp)
C
ChIP-peaks overlap in MOLM-13 cells
N-MLL1
MLL-AF9-HA
IRF8
SPI1
GFI1
D
ChIP-peaks overlap in THP1 cells
N-MLL1
C-AF9
IRF8
SPI1
GFI1
E
N-MLL1
C-AF9
IRF8
SPI1
GFI1
CEBPA
H3K4me3
H3K27ac
HOXA1 HOXA3 HOXA7 HOXA9 HOXA10 HOXA13
JAK2
F
MLL-AF9-HA
MLL-AF9-HA
N-MLL1
N-MLL1
GFI1
SPI1
IRF8
H3K27ac
HOXA3 HOXA7 HOXA9 HOXA11 HOXA13 HOTTIP
G
MLL-AF9
MBM ATH SNL CxxC AF9
breakpoint
M1 M2 M3 M4 M5
H
Input:
IB: Flag
M3-HA
M4-HA
M2-HA
M1-HA
M5-HA
IRF8-HA
SPI1-HA
IP: Flag
IB: HA
IRF8-HA
SPI1-HA
I
Input:
IB:GFI1
IP: GFI1
IB: Flag
Input
αGFI1
M1-Flag
M2-Flag
M3-Flag
M4-Flag
M5-Flag

**Fig. S4 MLL-AF9 interacts with IRF8, SPI1 or GFI1 and co-occupy on the chromatin**

(**A**) De novo motif analysis identifying consensus sequences bound by indicated key promonocyte TFs, using HOMER on sequences from chromatin regions bound by MLL-AF9 in THP1 and MOLM-13 cells. (**B**) Metaplot (top) and density plot (bottom) illustrating ChIP-seq profiles for GFI1 and SPI1 in MOLM-13 cells. (**C**) Venn diagrams depicting the shared binding peaks of MLL-AF9 with IRF8, SPI1, and GFI1 (left) and the overlap among binding peaks of MLL-AF9, IRF8, SPI1, and GFI1 in MOLM-13 cells. (**D**) Venn diagrams depicting the overlap among binding peaks of MLL-AF9, IRF8, SPI1, and GFI1 in THP1 cells. The percentage of overlap (left) and without overlap (right) binding peaks account for MLL-AF9 binding peaks number was highlighted in red.

(**E**) ChIP-seq gene tracks revealing the chromatin binding of N-MLL1, C-AF9, IRF8, SPI1, GFI1 and CEBPA accompanied by epigenetic marks H3K27ac and H3K4me3 at indicated gene loci in THP1 cells. ChIP-seq data for N-MLL1, C-AF9, H3K27ac and H3K4me3 were retrieved from GSE79899, while data for GFI1, IRF8, and SPI1 were obtained from GSE123872. (**F**) ChIP-seq gene tracks revealing the chromatin association of MLL-AF9-HA, N-MLL1, GFI1, SPI1, and IRF8, as well as the epigenetic mark H3K27ac at indicated gene loci in MOLM-13 cells. Chip-seq data for MLL-AF9-HA, N-MLL1, and H3K27ac were sourced from GSE173599, and data for IRF8 were obtained from GSE157636. (**G**) Diagram showing the structural information of different domains and chopped fragments in MLL-AF9. MBM, high-affinity Menin-binding motif; ATH, AT-Hook1/2/3; SNL, nuclear-localization signal 1/2. (**H-I**) Co-immunoprecipitation analysis illustrating MLL-AF9's interaction with IRF8, SPI1, and GFI1 through its distinct domains.

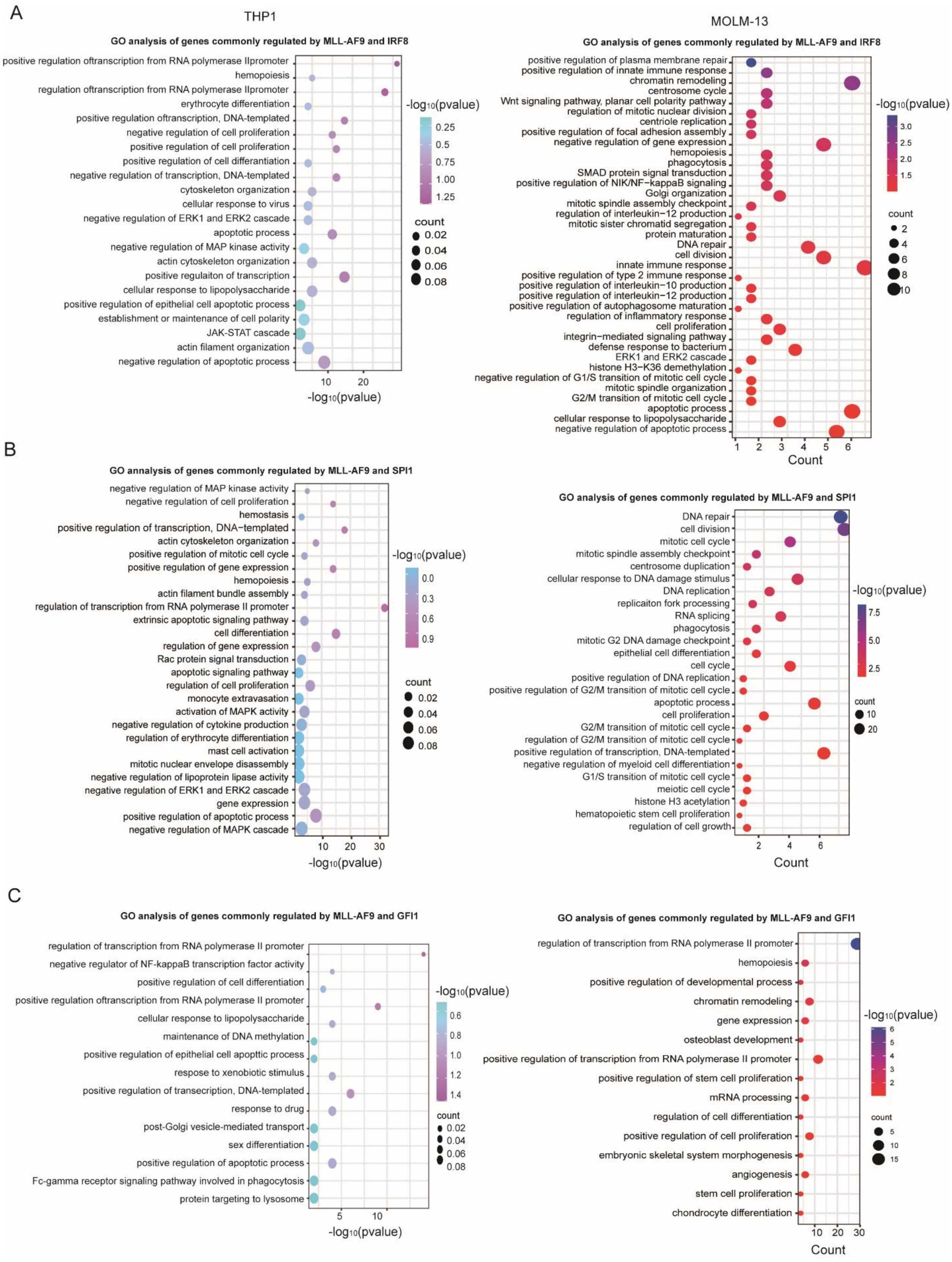

A
THP1
GO analysis of genes commonly regulated by MLL-AF9 and IRF8
positive regulation oftranscription from RNA polymerase IIpromoter
hemopoiesis
regulation oftranscription from RNA polymerase IIpromoter
erythrocyte differentiation
positive regulation oftranscription, DNA-templated
negative regulation of cell proliferation
positive regulation of cell proliferation
positive regulation of cell differantiation
negative regulation of transcription, DNA-templated
cytoskeleton organization
cellular response to virus
negative regulation of ERK1 and ERK2 cascade
apoptotic process
negative regulation of MAP kinase activity
actin cytoskeleton organization
positive regulaiton of transcription
cellular response to lipopolysaccharide
positive regulation of epithelial cell apoptotic process
establishment or maintenance of cell polarity
JAK-STAT cascade
actin filament organization
negative regulation of apoptotic process
10
20
-log10(pvalue)
-log10(pvalue)
0.25
0.5
0.75
1.00
1.25
count
0.02
0.04
0.06
0.08
MOLM-13
GO analysis of genes commonly regulated by MLL-AF9 and IRF8
positive regulation of plasma membrane repair
positive regulation of innate immune response
chromatin remodeling
centrosome cycle
Wnt signaling pathway, planar cell polarity pathway
regulation of mitotic nuclear division
centriole replication
positive regulation of focal adhesion assembly
negative regulation of gene expression
hemopoiesis
phagocytosis
SMAD protein signal transduction
positive regulation of NIK/NF−kappaB signaling
Golgi organization
mitotic spindle assembly checkpoint
regulation of interleukin−12 production
mitotic sister chromatid segregation
protein maturation
DNA repair
cell division
innate immune response
positive regulation of type 2 immune response
positive regulation of interleukin−10 production
positive regulation of interleukin−12 production
positive regulation of autophagosome maturation
regulation of inflammatory response
cell proliferation
integrin−mediated signaling pathway
defense response to bacterium
ERK1 and ERK2 cascade
histone H3−K36 demethylation
negative regulation of G1/S transition of mitotic cell cycle
mitotic spindle organization
G2/M transition of mitotic cell cycle
apoptotic process
cellular response to lipopolysaccharide
negative regulation of apoptotic process
1
2
3
4
5
6
Count
-log10(pvalue)
3.0
2.5
2.0
1.5
count
2
4
6
8
10
B
GO annalysis of genes commonly regulated by MLL-AF9 and SPI1
negative regulation of MAP kinase activity
negative regulation of cell proliferation
hemostasis
positive regulation of transcription, DNA−templated
actin cytoskeleton organization
positive regulation of mitotic cell cycle
positive regulation of gene expression
hemopoiesis
actin filament bundle assembly
regulation of transcription from RNA polymerase II promoter
extrinsic apoptotic signaling pathway
cell differentiation
regulation of gene expression
Rac protein signal transduction
apoptotic signaling pathway
regulation of cell proliferation
monocyte extravasation
activation of MAPK activity
negative regulation of cytokine production
regulation of erythrocyte differentiation
mast cell activation
mitotic nuclear envelope disassembly
negative regulation of lipoprotein lipase activity
negative regulation of ERK1 and ERK2 cascade
gene expression
positive regulation of apoptotic process
negative regulation of MAPK cascade
10
20
30
-log10(pvalue)
-log10(pvalue)
0.0
0.3
0.6
0.9
count
0.02
0.04
0.06
0.08
GO analysis of genes commonly regulated by MLL-AF9 and SPI1
DNA repair
cell division
mitotic cell cycle
mitotic spindle assembly checkpoint
centrosome duplication
cellular response to DNA damage stimulus
DNA replication
replicaiton fork processing
RNA splicing
phagocytosis
mitotic G2 DNA damage checkpoint
epithelial cell differentiation
cell cycle
positive regulation of DNA replication
positive regulation of G2/M transition of mitotic cell cycle
apoptotic process
cell proliferation
G2/M transition of mitotic cell cycle
regulation of G2/M transition of mitotic cell cycle
positive regulation of transcription, DNA-templated
negative regulation of myeloid cell differentiation
G1/S transition of mitotic cell cycle
meiotic cell cycle
histone H3 acetylation
hematopoietic stem cell proliferation
regulation of cell growth
2
4
6
Count
-log10(pvalue)
7.5
5.0
2.5
count
10
20
C
GO analysis of genes commonly regulated by MLL-AF9 and GFI1
regulation of transcription from RNA polymerase II promoter
negative regulator of NF-kappaB transcription factor activity
positive regulation of cell differentiation
positive regulation oftranscription from RNA polymerase II promoter
cellular response to lipopolysaccharide
maintenance of DNA methylation
positive regulation of epithelial cell apoptotic process
resposne to xenobiotic stimulus
positive regulation of transecription, DNA-templated
response to drug
post-Golgi vesicle-mediated transport
sex differentiation
positive regulation of apoptotic process
Fc-gamma receptor signaling pathway involved in phagocytosis
protein targeting to lysosome
5
10
-log10(pvalue)
-log10(pvalue)
0.6
0.8
1.0
1.2
1.4
count
0.02
0.04
0.06
0.08
GO analysis of genes commonly regulated by MLL-AF9 and GFI1
regulation of transcription from RNA polymerase II promoter
hemopoiesis
positive regulation of developmental process
chromatin remodeling
gene expression
osteoblast development
positive regulation of transcription from RNA polymerase II promoter
positive regulation of stem cell proliferation
mRNA processing
regulation of cell differentiation
positive regulation of cell proliferation
embryonic skeletal system morphogenesis
angiogenesis
stem cell proliferation
chondrocyte differentiation
10
20
30
Count
-log10(pvalue)
6
5
4
3
2
count
5
10
15

**Fig. S5 GO analysis of genes commonly regulated by MLL-AF9 and key promonocyte TFs in THP1 and MOLM-13 cells**

(**A-C**) GO analysis of genes commonly regulated by MLL-AF9 alongside IRF8 (A), SPI1 (B), or GFI1 (C) in THP1 (left) and MOLM-13 (right) cells.

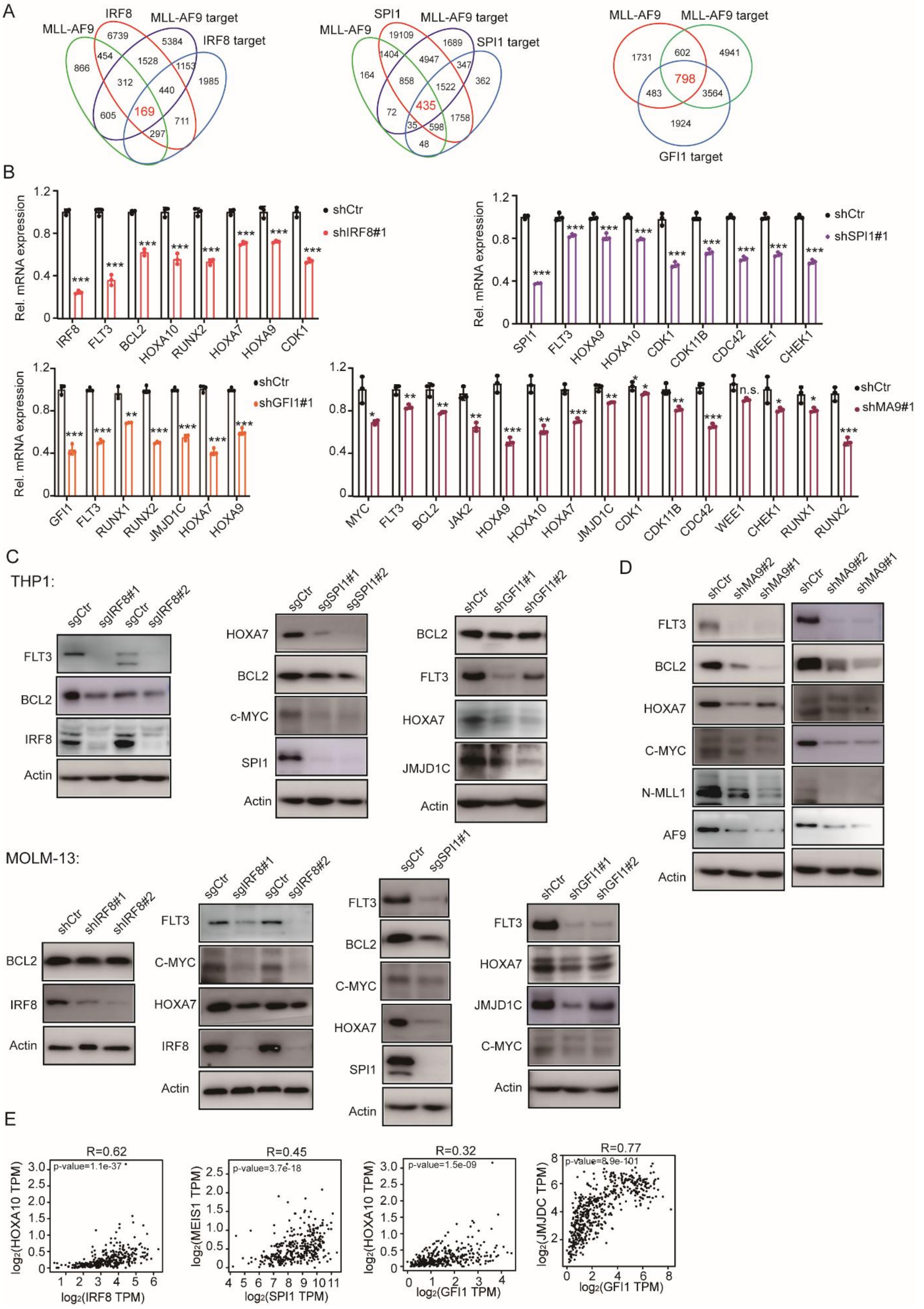

A
MLL-AF9
IRF8
MLL-AF9 target
IRF8 target
169
SPI1
SPI1 target
435
GFI1 target
798
B
Rel. mRNA expression
shCtr
shIRF8#1
shSPI1#1
shGFI1#1
shMA9#1
C
THP1:
MOLM-13:
D
E
R=0.62
p-value=1.1e-37
log2(HOXA10 TPM)
log2(IRF8 TPM)
R=0.45
p-value=3.7e-18
log2(MEIS1 TPM)
log2(SPI1 TPM)
R=0.32
p-value=1.5e-09
log2(GFI1 TPM)
R=0.77
p-value=8.9e-101
log2(JMJDC TPM)

**Fig. S6 MLL-AF9 and key promonocyte TFs co-regulated the expression of genes crucial for AML hallmarks**

(**A**) Venn diagrams indicating the overlap between genes bound by the indicated proteins and those downregulated following knockdown of specified genes in MOLM-13 cells. The number of common downregulated and co-bound genes between MLL-AF9 and IRF8, SPI1 or GFI1 was highlighted in red. (**B**) RT-qPCR analysis revealing genes commonly downregulated in MOLM-13 cells upon knockdown of MLL-AF9 (with shMA9#1) and TFs IRF8 (with shIRF8#1), SPI1 (with shSPI1#1) or GFI1 (with shGFI1#1). (**C-D**) Western-blot results revealing the expression levels of indicated genes in THP1 (b) and MOLM-13 (c) cells following knockout of IRF8 (with sgIRF8#1 and sgIRF8#2) or SPI1 (with sgSPI1#1 and sgSPI1#2), or knockdown of IRF8 (with shIRF8#1 and shIRF8#2), GFI1 (with shGFI1#1 and shGFI1#2) or MLL-AF9 (with shMA9#1 and shMA9#2). Control sgRNA (sgCtr) or shRNA (shCtr) transduced cells serve as controls. (**E**) Dot plots illustrating the positive expression correlation between IRF8, SPI1 or GFI1 with genes key for AML hallmarks in AML cells, analyzed using GEPIA2.

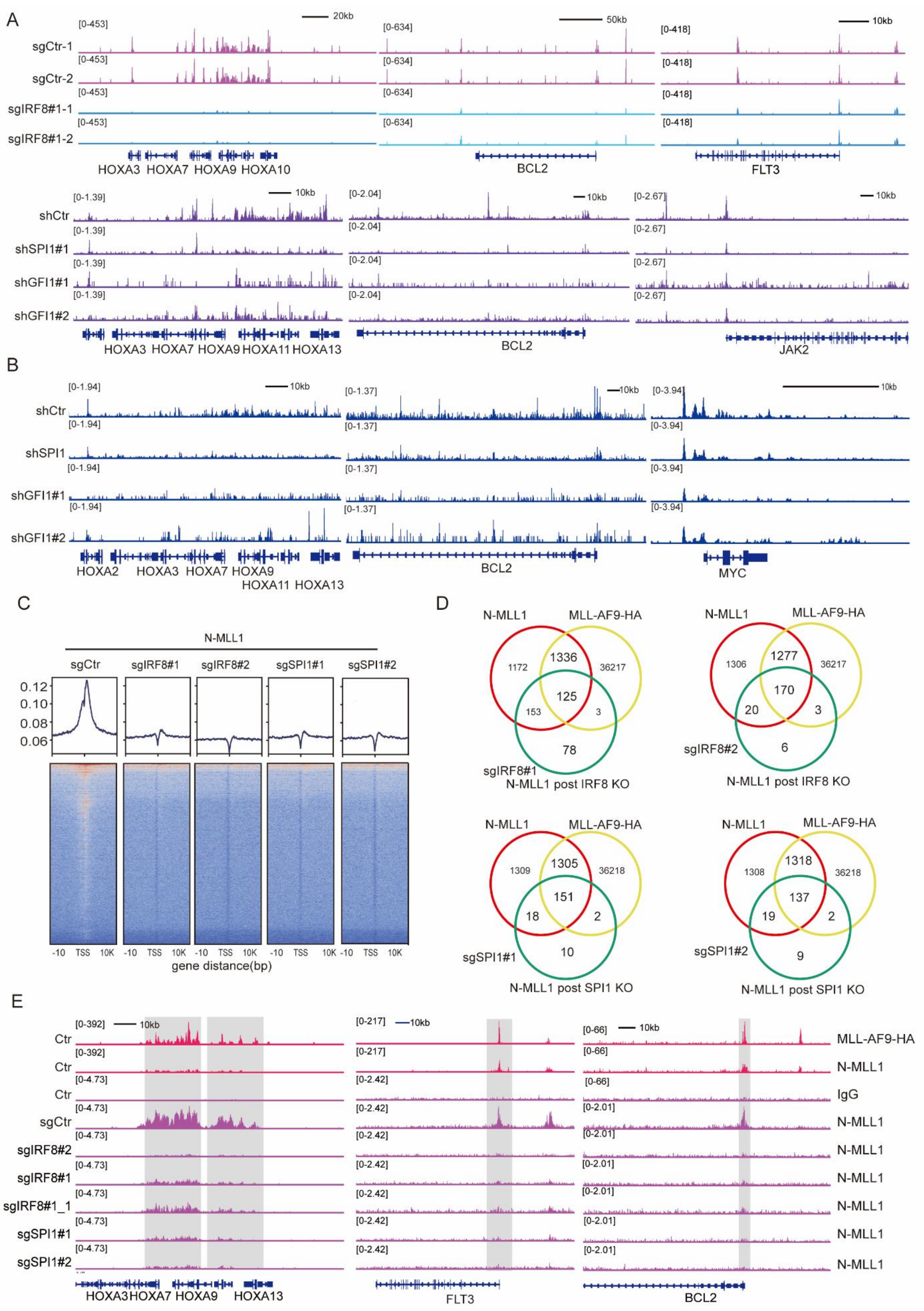
A
sgCtr-1
sgCtr-2
sgIRF8#1-1
sgIRF8#1-2
HOXA3 HOXA7 HOXA9 HOXA10
BCL2
FLT3
shCtr
shSPI1#1
shGFI1#1
shGFI1#2
HOXA3 HOXA7 HOXA9 HOXA11 HOXA13
BCL2
JAK2
B
shCtr
shSPI1
shGFI1#1
shGFI1#2
HOXA2 HOXA3 HOXA7 HOXA9
HOXA11 HOXA13
BCL2
MYC
C
N-MLL1
sgCtr
sgIRF8#1
sgIRF8#2
sgSPI1#1
sgSPI1#2
gene distance(bp)
D
N-MLL1
MLL-AF9-HA
sgIRF8#1
N-MLL1 post IRF8 KO
sgIRF8#2
sgSPI1#1
N-MLL1 post SPI1 KO
sgSPI1#2
E
Ctr
sgCtr
sgIRF8#2
sgIRF8#1
sgIRF8#1_1
sgSPI1#1
sgSPI1#2
HOXA3 HOXA7 HOXA9 HOXA13
FLT3
BCL2
MLL-AF9-HA
N-MLL1
IgG

**Fig. S7 Disruption of reinforced promonocyte identity reduces chromatin accessibility and MLL-AF9 chromatin binding at genes essential for AML hallmarks**

(**A**) ATAC-seq gene tracks showing changes in chromatin accessibility at HOXA family, BCL2 and FLT3 loci in THP1 cells following IRF8 knockout (with sgIRF8#1). Cells transduced with control sgRNAs (sgCtr) serve as controls. ATAC-seq gene tracks depicting chromatin accessibility changes at HOXA family, BCL2, and JAK2, loci in THP1 cells post SPI1 or GFI1 knockdown (with shSPI1#1, shGFI1#1, or shGFI1#2 respectively). Cells transduced with control shRNA (shCtr) serve as controls. (**B**) ATAC-seq gene tracks displaying chromatin accessibility at HOXA family, BCL2, and MYC loci in MOLM-13 cells after SPI1 or GFI1 knockdown (with shSPI1, shGFI1#1, or shGFI1#2 respectively). Cells transduced with control shRNA (shCtr) serve as controls. (**C**) Metaplot (top) and density (bottom) plots depicting ChIP-seq profiles for N-MLL1 in MOLM-13 cells transduced with sgRNAs targeting IRF8 (sgIRF8#1 and sgIRF8#2) or SPI1 (sgSPI1#1 and sgSPI1#2). Cells transduced with control sgRNAs (sgCtr) serve as controls. (**D**) Venn diagrams showing the overlap in MOLM-13 cells among binding peaks of N-MLL1, and MLL-AF9-HA, and those lost upon knockout of IRF8 (using sgIRF8#1 or sgIRF8#2) or SPI1 (using sgSPI1#1 or sgSPI1#2). ChIP-seq data for MLL-AF9-HA were sourced from GSE173599. (**E**) ChIP-seq gene tracts displaying MLL-AF9 binding peaks at specified gene loci in MOLM-13 expressing control (sgCtr), IRF8 (sgIRF8#1 and sgIRF8#2), or SPI1 (sgSPI1#1 and sgSPI1#2) sgRNAs. ChIP-seq data for MLL-AF9-HA were sourced from GSE173599.

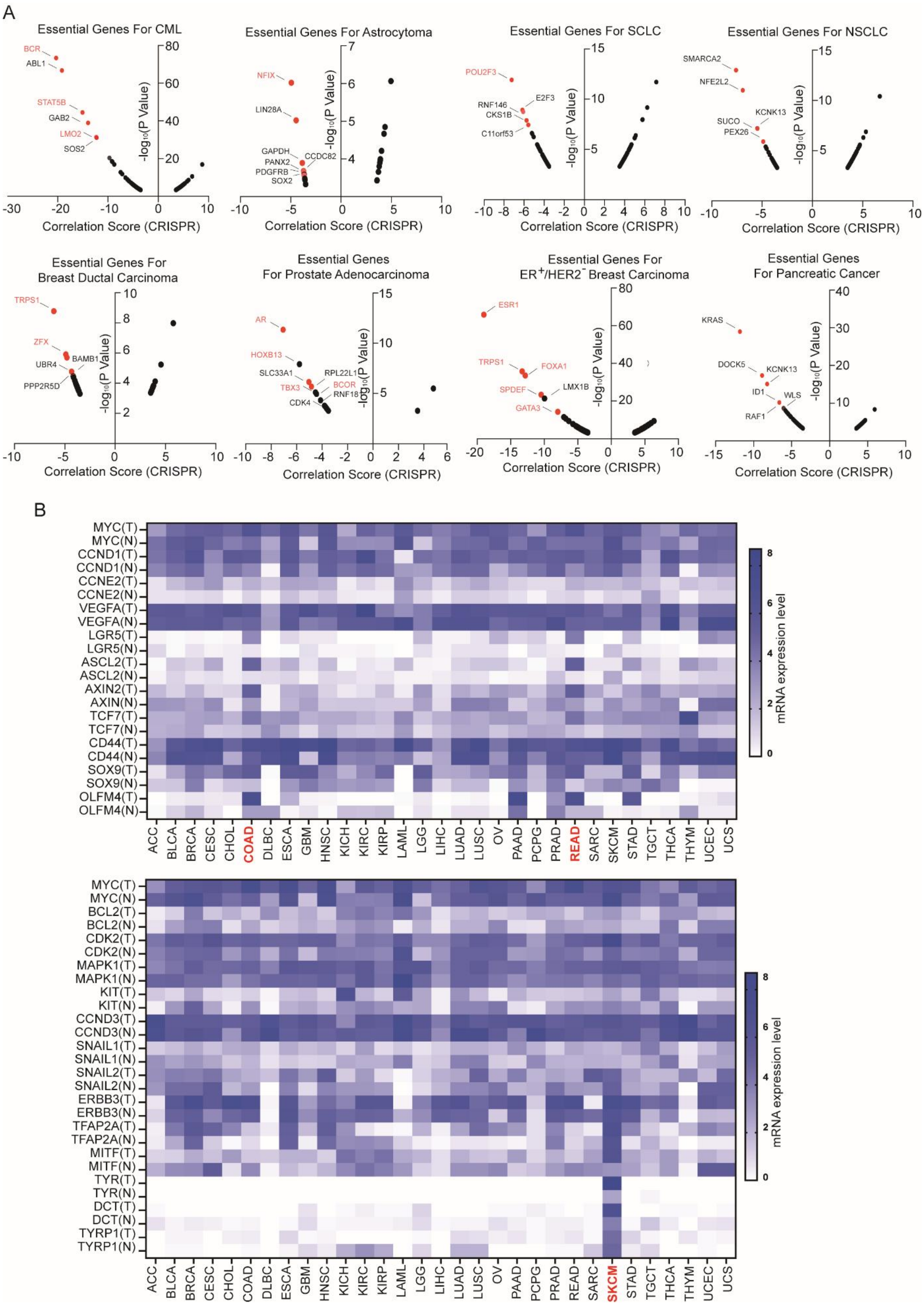
A
Essential Genes For CML
Essential Genes For Astrocytoma
Essential Genes For SCLC
Essential Genes For NSCLC
Essential Genes For Breast Ductal Carcinoma
Essential Genes For Prostate Adenocarcinoma
Essential Genes For ER+/HER2- Breast Carcinoma
Essential Genes For Pancreatic Cancer
Correlation Score (CRISPR)
-log10(P Value)
BCR
ABL1
STAT5B
GAB2
LMO2
SOS2
NFIX
LIN28A
GAPDH
PANX2
CCDC82
PDGFRB
SOX2
POU2F3
RNF146
E2F3
CKS1B
C11orf53
SMARCA2
NFE2L2
SUCO
KCNK13
PEX26
TRPS1
ZFX
BAMB1
UBR4
PPP2R5D
AR
HOXB13
SLC33A1
RPL22L1
TBX3
BCOR
RNF18
CDK4
ESR1
TRPS1
FOXA1
SPDEF
LMX1B
GATA3
KRAS
DOCK5
KCNK13
ID1
WLS
RAF1
B
mRNA expression level
MYC(T)
MYC(N)
CCND1(T)
CCND1(N)
CCNE2(T)
CCNE2(N)
VEGFA(T)
VEGFA(N)
LGR5(T)
LGR5(N)
ASCL2(T)
ASCL2(N)
AXIN2(T)
AXIN(N)
TCF7(T)
TCF7(N)
CD44(T)
CD44(N)
SOX9(T)
SOX9(N)
OLFM4(T)
OLFM4(N)
BCL2(T)
BCL2(N)
CDK2(T)
CDK2(N)
MAPK1(T)
MAPK1(N)
KIT(T)
KIT(N)
CCND3(T)
CCND3(N)
SNAIL1(T)
SNAIL1(N)
SNAIL2(T)
SNAIL2(N)
ERBB3(T)
ERBB3(N)
TFAP2A(T)
TFAP2A(N)
MITF(T)
MITF(N)
TYR(T)
TYR(N)
DCT(T)
DCT(N)
TYRP1(T)
TYRP1(N)
ACC BLCA BRCA CESC CHOL COAD DLBC ESCA GBM HNSC KICH KIRC KIRP LAML LGG LIHC LUAD LUSC OV PAAD PCPG PRAD READ SARC SKCM STAD TGCT THCA THYM UCEC UCS

**Fig. S8 The dependency of intrinsic cell identities on the maintenance of a broad spectrum of cancer types**

(**A**) Graphic representations depicting the correlation between the knockout of specific genes and the survival of multiple cancer types. Identity genes were highlighted in red. (**B**) Heatmap depicting the differential expression of genes vital for intestinal stem cell traits and cancer hallmarks in colorectal cancer (top) and genes crucial both neural crest cell traits and cancer hallmarks in melanoma (bottom), comparing their respective normal tissue counterparts. ACC, Adrenocortical carcinoma; BLCA, Bladder Urothelial Carcinoma; BRCA, Breast invasive carcinoma; CESC, Cervical squamous cell carcinoma and endocervical adenocarcinoma; CHOL, Cholangiocarcinoma; COAD, Colon adenocarcinoma; DLBC, Lymphoid Neoplasm Diffuse Large B-cell Lymphoma; ESCA, Esophageal carcinoma; GBM, Glioblastoma multiforme; HNSC, Head and Neck squamous cell carcinoma; KICH, Kidney Chromophobe; KIRC, Kidney renal clear cell carcinoma; KIRP, Kidney renal papillary cell carcinoma; LAML, Acute Myeloid Leukemia; LGG, Brain Lower Grade Glioma; LIHC, Liver hepatocellular carcinoma; LUAD, Lung adenocarcinoma; OV, Ovarian serous cystadenocarcinoma; LUSC, Lung squamous cell carcinoma; PAAD, Pancreatic adenocarcinoma; PCPG, Pheochromocytoma and Paraganglioma; PRAD,Prostate adenocarcinoma; READ, Rectum adenocarcinoma; SARC, Sarcoma; SKCM, Skin Cutaneous Melanoma; STAD, Stomach adenocarcinoma; TGCT, Testicular Germ Cell Tumors; THCA, Thyroid carcinoma; THYM, Thymoma; UCEC, Uterine Corpus Endometrial Carcinoma; UCS, Uterine Carcinosarcoma.

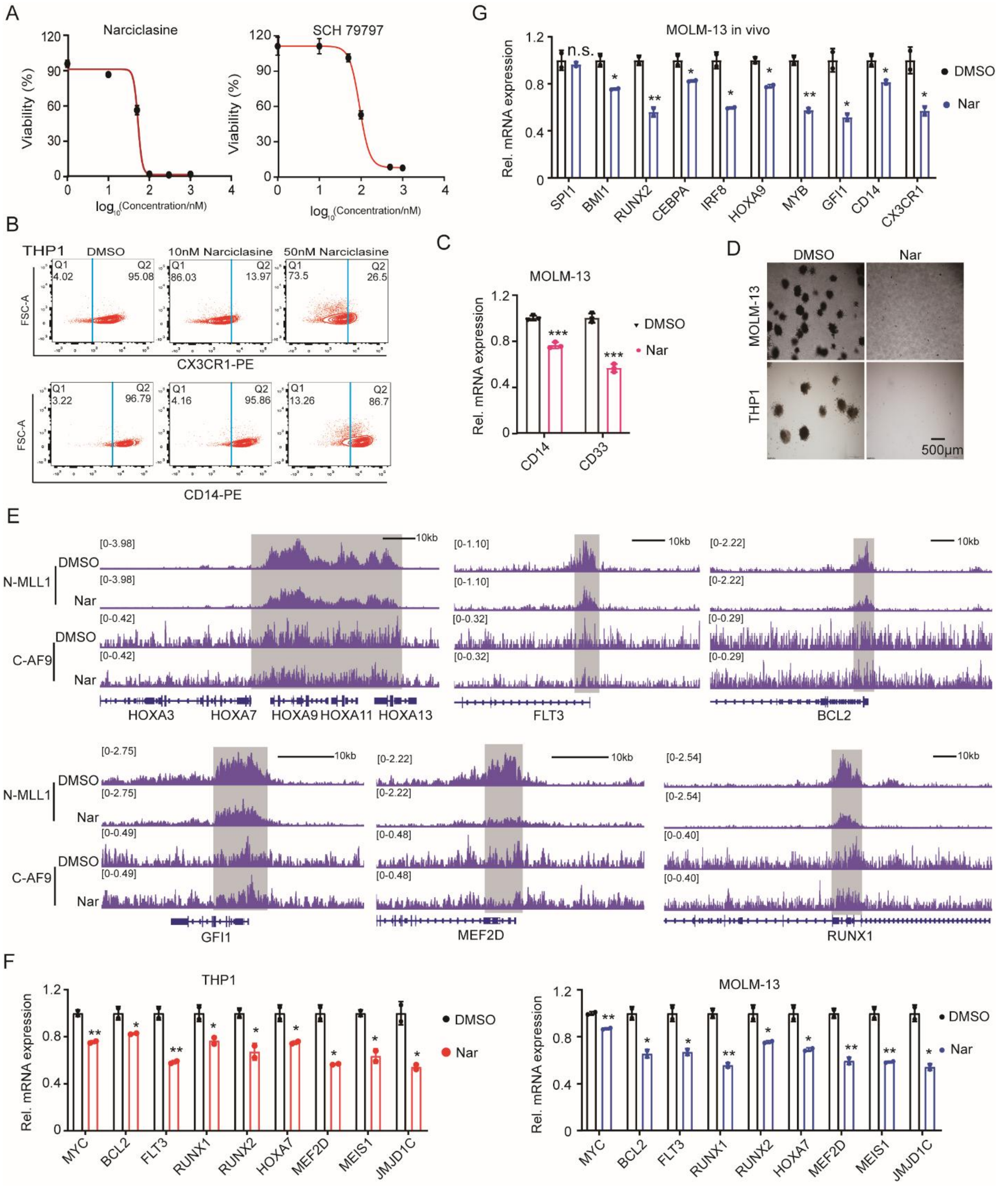

**Fig. S9 Narciclasine inhibits cancer hallmarks, disrupts promonocyte identity and decreases MLL-AF9 occupancy on the chromatin in AML-M5$^{MLL\text{-}AF9}$ cells**

(**A**) Dose-response curves showing the inhibitory effects on the number of MOLM-13 cells treated with indicated concentrations of narciclasine and SCH 79797 for 36h. (**B**) Flow cytometry analysis depicting CX3CR1 (top) and CD14 (bottom) expression in THP1 cells following treatment with DMSO, or 10nM and 50nM narciclasine for 48h. (**C**) RT-qPCR results displaying differential expression levels of CD14 and CD33 mRNA in MOLM-13 cells after treatment with 50 nM narciclasine (Nar) for 12h. (**D**) Bright-field images revealing colony formation in methylcellulose culture of THP1 and MOLM-13 cells treated with DMSO or 50nM narciclasine for 7 days. (**E**) Chip-seq gene tracks showing MLL-AF9 binding peaks at selected loci in THP1 cells treated with DMSO or 50nM narciclasine (Nar) for 12h. (**F**) RT-qPCR results displaying the expression levels of genes pivotal for promonocyte traits and AML hallmarks in THP1 and MOLM-13 cells after treatment with DMSO or 50 nM narciclasine (Nar) for 12h. (**G**) RT–qPCR results showing the expression levels of key promonocyte TFs and surface markers in human $CD45^+$ leukemia cells isolated from bone marrow of NSG mice. NSG mice were transplanted with MOLM-13 cells, and then treated with vehicle (0.001% DMSO) or Narciclasine (Nar) every other day for 8 days.

**Table S1 shRNA and sgRNA sequences**

| Primer Name | Sequence (5'to3') |
|---|---|
| IRF8-SgRNA1-F | caccgATTGACAGTAGCATGTATCC |
| IRF8-SgRNA1-R | aaacGGATACATGCTACTGTCAATc |
| IRF8-SgRNA2-F | caccgAGAGCATGTTCCGGATCCCT |
| IRF8-SgRNA2-R | aaacAGGGATCCGGAACATGCTCTc |
| SPI1-SgRNA1-F | caccgAGACCTGGTGCCCTATGACA |
| SPI1-SgRNA1-R | aaacTGTCATAGGGCACCAGGTCTc |
| SPI1-SgRNA2-F | caccgGCACGTCCTCGATACCCCCA |
| SPI1-SgRNA2-R | aaacTGGGGGTATCGAGGACGTGCc |
| GFI1-SgRNA1-F | caccgGTTGGGGTAACCCACCCGAT |
| GFI1-SgRNA1-R | aaacATCGGGTGGGTTACCCCAACc |
| GFI1-SgRNA2-F | caccgCCGGTACATTCTCTAAACGG |
| GFI1-SgRNA2-R | aaacCCGTTTAGAGAATGTACCGGc |
| IRF8-shRNA#1 | CCGGCCATACAAAGTTTACCGAATTCTCGAGAATTCGGTAAACTTTGTATGGTTTTT |
| IRF8-shRNA#2 | CCGGGCTTTGAATAAGAGCCCAGATCTCGAGATCTGGGCTCTTATTCAAAGCTTTTT |
| IRF8-shRNA#3 | CCGGGCCCGCATCATGATTAAAGAACTCGAGTTCTTTAATCATGATGCGGGCTTTTT |
| SPI1-shRNA#1 | CCGGGCCCTATGACACGGATCTATACTCGAGTATAGATCCGTGTCATAGGGCTTTTT |
| SPI1-shRNA#2 | CCGGCCTCCACATCCCGCTTCGCCTCTCGAGAGGCGAAGCGGGATGTGGAGGTTTTT |
| SPI1-shRNA#3 | CCGGCGGATCTATACCAACGCCAAACTCGAGTTTGGCGTTGGTATAGATCCGTTTTT |
| GFI1-shRNA#1 | CCGGTGCCTTTCAAACCGTACTCATCTCGAGATGAGTACGGTTTGAAAGGCATTTTT |
| GFI1-shRNA#2 | CCGGCATCAAGTGCAGCAAGGTGTTCTCGAGAACACCTTGCTGCACTTGATGTTTTT |
| GFI1-shRNA#3 | CCGGCGACCTCTGTGGGAAGGGTTTCTCGAGAAACCCTTCCCACAGAGGTCGTTTTT |

**Table S2 MLL-AF9 binds loci near promonocyte TFs**

| Item | Genome loci |
|---|---|
| IRF8-P1 | Chr16 dna:chromosome chromosome:GRCh38:16:85898500:85900500:1 |
| IRF8-E1 | Chr16 dna:chromosome chromosome:GRCh38:16:85982200:85983200:1 |
| IRF8-E2 | Chr16 dna:chromosome chromosome:GRCh38:16:85983250:85984750:1 |
| IRF8-E3 | Chr16 dna:chromosome chromosome:GRCh38:16:85984750:85985750:1 |
| SPI1-P1 | Chr11 dna:chromosome chromosome:GRCh38:11:47376500:47378000:1 |
| SPI1-P2 | Chr11 dna:chromosome chromosome:GRCh38:11:47377500:47379000:1 |
| SPI1-P3 | Chr11 dna:chromosome chromosome:GRCh38:11:47390500:47392000:1 |
| SPI1-P4 | Chr11 dna:chromosome chromosome:GRCh38:11:47392500:47393750:1 |
| SPI1-P5 | Chr11 dna:chromosome chromosome:GRCh38:11:47393750:47395500:1 |
| GFI1-P1 | Chr1 dna:chromosome chromosome:GRCh38:1:92481000:92482000:1 |
| GFI1-P2 | Chr1 dna:chromosome chromosome:GRCh38:1:92482000:92484000:1 |
| GFI1-P3 | Chr1 dna:chromosome chromosome:GRCh38:1:92484000:92486000:1 |
| GFI1-P4 | Chr1 dna:chromosome chromosome:GRCh38:1:92486000:92488000:1 |
| CEBPA-P1 | Chr19 dna:chromosome chromosome:GRCh38:19:33298500:33300000:1 |
| CEBPA-P2 | Chr19 dna:chromosome chromosome:GRCh38:19:33300000:33301600:1 |
| CEBPA-P3 | 19 dna:chromosome chromosome:GRCh37:19:33301000:33302700:1 |

**Table S3. RT-qPCR primer sequences**

| Primer Name | Sequence (5'to3') |
|---|---|
| IRF8-F | AGGTCTTCGACACCAGCCAGTT |
| IRF8-R | GCACGAGAATGAGTTTGGAGCG |
| SPI1-F | GACACGGATCTATACCAACGCC |
| SPI1-R | CCGTGAAGTTGTTCTCGGCGAA |
| GFI1-F | GCTTCAAGAGGTCATCCACACTG |
| GFI1-R | ACCTGGCACTTGTGAGGCTTCT |
| ACTB-F | CACCATTGGCAATGAGCGGTTC |
| ACTB-R | AGGTCTTTGCGGATGTCCACGT |
| MYB-F | CAGTTCGCAGACCTCCTGTTGA |
| MYB-R | TCCAGCTCCTTCAGAGTCTGCA |
| LMO2-F | GCGCCTCTACTACAAACTGGGC |
| LMO2-R | CTCATAGGCACGAATCCGCTTG |
| RUNX1-F | CCACCTACCACAGAGCCATCAA |
| RUNX1-R | TTCACTGAGCCGCTCGGAAAAG |
| RUNX2-F | CCCAGTATGAGAGTAGGTGTCC |
| RUNX2-R | GGGTAAGACTGGTCATAGGACC |
| CD11b-F | CAGACAGGAAGTAGCAGCTCCT |
| CD11b-R | CTGGTCATGTTGATGAAGGTGCT |
| CD14 -F | GACCTAAAGATAACCGGCACC |
| CD14 -R | GCAATGCTCAGTACCTTGAGG |
| CSF3R -F | TCAAGTTGGTGCTATGGCAAGG |
| CSF3R -R | GCTCCCAGTCTCCACAGAATC |
| CSF1R-F | GGGAATCCCAGTGATAGAGCC |
| CSF1R-R | TTGGAAGGTAGCGTTGTTGGT |
| CD16-F | GGTGACTTGTCCACTCCAGTGT |
| CD16 -R | ACCATTGAGGCTCCAGGAACAC |
| CX3CR1-F | CACAAAGGAGCAGGCATGGAAG |
| CX3CR1-R | CAGGTTCTCTGTAGACACAAGGC |
| CD13-F | GCTGTTTGACGCCATCTCCTAC |
| CD13-R | GTTCTGGTAGGCAAAGGTGTGG |
| CD33-F | GTGACTACGGAGAGAACCATCC |
| CD33-R | GCTGTAACACCAGCTCCTCCAA |
| CD34-F | CCTCAGTGTCTACTGCTGGTCT |
| CD34-R | GGAATAGCTCTGGTGGCTTGCA |
| CD64-F | ATACAGGTGCCAGAGAGGTCTC |
| CD64-R | CCAGCTTATCCTTCCACGCATG |
| CD4-F | CCTCCTGCTTTTCATTGGGCTAG |
| CD4-R | TGAGGACACTGGCAGGTCTTCT |
| CD45-F | CTTCAGTGGTCCCATTGTGGTG |
| CD45-R | CCACTTTGTTCTCGGCTTCCAG |
| CD15/FUT4-F | GGGTTTGGATGAACTTCGAGTCG |
| CD15/FUT4-R | GGTAGCCATAAGGCACAAAGACG |
| c Kit-F | CACCGAAGGAGGCACTTACACA |

| | |
|---|---|
| c Kit-R | TGCCATTCACGAGCCTGTCGTA |
| CD36-F | CAGGTCAACCTATTGGTCAAGCC |
| CD36-R | GCCTTCTCATCACCAATGGTCC |
| CD11c-F | GATGCTCAGAGATACTTCACGGC |
| CD11c-R | CCACACCATCACTTCTGCGTTC |
| CD18-F | AGTCACCTACGACTCCTTCTGC |
| CD18-R | CAAACGACTGCTCCTGGATGCA |
| CD35-F | TAGGTGTCAGCCTGGCTTTGTC |
| CD35-R | GACATCTGGAGGTGGCTGACAT |
| MRC1-F | AGCCAACACCAGCTCCTCAAGA |
| MRC1-R | CAAAACGCTCGCGCATTGTCCA |
| CCR1-F | CAACTCCGTGCCAGAAGGTGAA |
| CCR1-R | GTTCAGGAGGTAGATGCTGGTC |
| CCR5-F | TCTCTTCTGGGCTCCCTACAAC |
| CCR5-R | CCAAGAGTCTCTGTCACCTGCA |
| qBCL2-F | ATCGCCCTGTGGATGACTGAGT |
| qBCL2-R | GCCAGGAGAAATCAAACAGAGGC |
| qFLT3-F | AGACTGTCGCTGGGTCCAAGAT |
| qFLT3-R | GGAGATGTTGGTCTGGACGAAG |
| qJAK2-F | CCAGATGGAAACTGTTCGCTCAG |
| qJAK2-R | GAGGTTGGTACATCAGAAACACC |
| qMEF2D-F1 | CCAGCGAATCACCGACGAG |
| qMEF2D-R1 | GCAGTCACATAGCACGCTC |
| qMEF2D-F2 | CGTGCTATGTGACTGCGAGAT |
| qMEF2D-R2 | GCGTCGGTACTTGTCCTCC |
| qHOXA3-F | CCTGCTCAACTCACCCACAGTG |
| qHOXA3-R | TCTTGTCGCCAGCGCAGCTTTC |
| qHOXA5-F | AACCCCAGATCTACCCCTGGAT |
| qHOXA5-R | CAGGGTCTGGTAGCGCGTGTA |
| qHOXA7-F | GCTGAGGCCAATTTCCGCATCT |
| qHOXA7-R | GTAGCGGTTGAAGTGGAACTCC |
| qHOXA10-F | CTTCCGAGAGCAGCAAAGCCTC |
| qHOXA10-R | TCCAGTGTCTGGTGCTTCGTGT |
| qJMJD1C-F | TCCTGTCAGACCTTCCAGTGCA |
| qJMJD1C-R | GTGGATGCAACAGACCGTAATGG |
| qc-Myc-F | CCTGGTGCTCCATGAGGAGAC |
| qc-Myc-R | CAGACTCTGACCTTTTGCCAGG |
| qPUF60-F | CTTGGAGCAGATGAACTCGGTG |
| qPUF60-R | CTCCTCAGCCAACTGGTCTATG |
| qHOXA11-F | CAGCAGAGGAGAAAGAGCGGC |
| qHOXA11-R | TCGGATCTGGTACTTGGTATAGG |
| qIRF2BP2-F | CGAGAGCAAGTTTAAGAAGGAGC |
| qIRF2BP2-R | TGGTTCTGGAGAGGGCTTCCTT |
| qMED13L-F | GAAGTCCCAAGCCCGAGGAAAT |
| qMED13L-R | CTCGGCAACATCTTCAGTGGAG |
